# The interactome of the Bakers’ yeast peroxiredoxin Tsa1 implicates it in the redox regulation of intermediary metabolism, glycolysis and zinc homeostasis

**DOI:** 10.1101/2025.02.18.638137

**Authors:** Colin W. MacDiarmid, Janet Taggart, Yirong Wang, Ajay Vashisht, Xin Qing, James A. Wohlschlegel, David J. Eide

## Abstract

Zinc (Zn) is an essential nutrient supporting a range of critical processes. In the yeast *Saccharomyces cerevisiae*, Zn deficiency induces a transcriptional response mediated by the Zap1 activator, which controls a regulon of ∼80 genes. A subset support zinc homeostasis by promoting zinc uptake and its distribution between compartments, while the remainder mediate an “adaptive response” to enhance fitness of zinc deficient cells. The peroxiredoxin Tsa1 is a Zap1-regulated adaptive factor essential for the growth of Zn deficient cells. Tsa1 can function as an antioxidant peroxidase, protein chaperone, or redox sensor: the latter activity oxidizes associated proteins via a redox relay mechanism. We previously reported that in Zn deficient cells, Tsa1 inhibits pyruvate kinase (Pyk1) to conserve phosphoenolpyruvate for aromatic amino acid synthesis. However, this regulation makes a relatively minor contribution to fitness in low zinc, suggesting that Tsa1 targets other pathways important to adaptation. Consistent with this model, the redox sensor function of Tsa1 was essential for growth of ZnD cells. Using an MBP-tagged version of Tsa1, we identified a redox-sensitive non-covalent interaction with Pyk1, and applied this system to identify multiple novel interacting partners. This interactome implicates Tsa1 in the regulation of critical processes including many Zn-dependent metabolic pathways. Interestingly, Zap1 was a preferred Tsa1 target, as Tsa1 strongly promoted the oxidation of Zap1 activation domain 2, and was essential for full Zap1 activity. Our findings reveal a novel posttranslational response to Zn deficiency, overlain on and interconnected with the Zap1-mediated transcriptional response.

**Graphical abstract:** In ZnD cells, Tsa1 mediates metabolic adaptation by regulation of glycolytic enzymes Pyk1 and Fba1, and supports activity of the Zap1 transcriptional activator via oxidation of the AD2 domain.

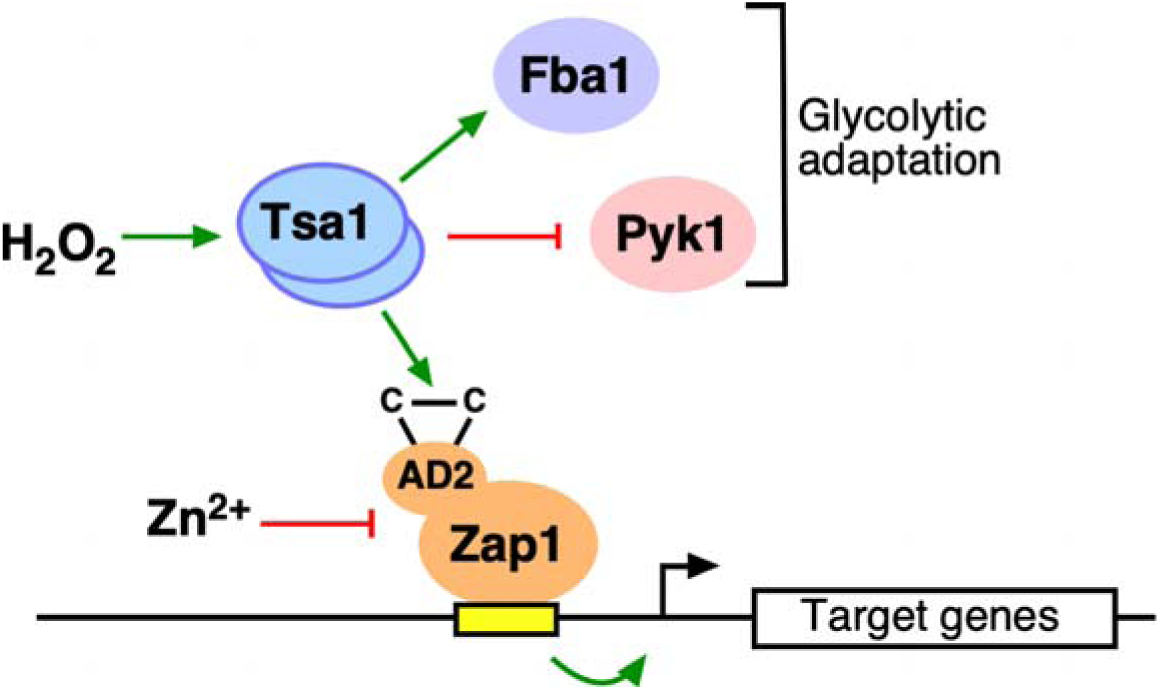

## INTRODUCTION

Zinc (Zn) is essential for all organisms, and cells must tightly regulate its accumulation to support essential functions while avoiding toxicity [1]. Zn deficiency is a common human health issue, of particular prevalence in infants, pregnant women and the elderly [2]. Additionally, Zn availability and homeostasis is critical to both microbial virulence and the host immune response [3, 4]. Among eukaryotes, mechanisms of Zn homeostasis are arguably best understood in the yeast *Saccharomyces cerevisiae* [5–14]. Approximately 10% of yeast proteins require Zn [15], and to manage this demand, the Zn-responsive activator Zap1 transcriptionally regulates ∼80 genes [16, 17]. Zap1-regulated genes fall into two broad classes, those mediating Zn homeostasis (*e.g.,* transporters) [10–14, 18, 19], and those that facilitate adaptation to Zn deficiency without directly influencing zinc availability [8, 9, 17, 20].

One such Zap1-regulated gene encodes one of the most critical adaptive factors, the peroxiredoxin (PR) Tsa1 [9, 20, 21]. Loss of Tsa1 function causes a severe growth defect in zinc-deficient (ZnD) cells. PR’s are ubiquitous and abundant proteins implicated in human disease [22–28] and have three well described functions: first, their thioredoxin-dependent peroxidase activity [29] reduces H_2_O_2_, lipid hydroperoxides, and some reactive nitrogen species. Tsa1 is a member of the 2-Cys subclass of PR’s that form obligate homodimers. During the catalytic cycle, the peroxidatic cysteine of one monomer (C48 of Tsa1) is oxidized by substrate to a sulfenic acid, which then reacts with the resolving cysteine of the partner monomer (C171). The resulting disulfide-linked dimers can be reduced by thioredoxin. The high affinity and abundance of thiol peroxidases in most cells forms their primary defense against ROS and RNS accumulation. The second known function of PR enzymes is exhibited by the so-called “sensitive” group of 2-Cys PR’s, including Tsa1. Exposure of these enzymes to excess substrate hyperoxidizes the peroxidatic cysteine, inactivating peroxidase function but activating a holdase-type protein chaperone [30–35]. Although the physiological role of this activity is not well understood, it has been suggested to stabilize the apoprotein forms of zinc-dependent proteins that accumulate in ZnD cells [9], and also to counteract the aggregation of misfolded proteins during yeast aging [36]. Finally, some PR enzymes can act as redox sensors [37–41]. In this role, the peroxidatic cysteine is oxidized to a sulfenic acid, which reacts with a cysteine on an associated target to form a mixed disulfide intermediate. Disulfide exchange with another target cysteine results in the introduction of a disulfide bond in the target, which can modulate its activity [40–48] (**Fig. 1A**). Such “redox relays” were originally described in yeast, where Tsa1 and Orp1 enable the transcriptional response to ROS by promoting oxidation of the transcriptional activator Yap1 [41, 46, 47, 49–51]. Subsequent studies have identified PR-mediated relays in microbial, plant and vertebrate systems [38, 39, 43, 44, 52–56].

**Figure 1.**
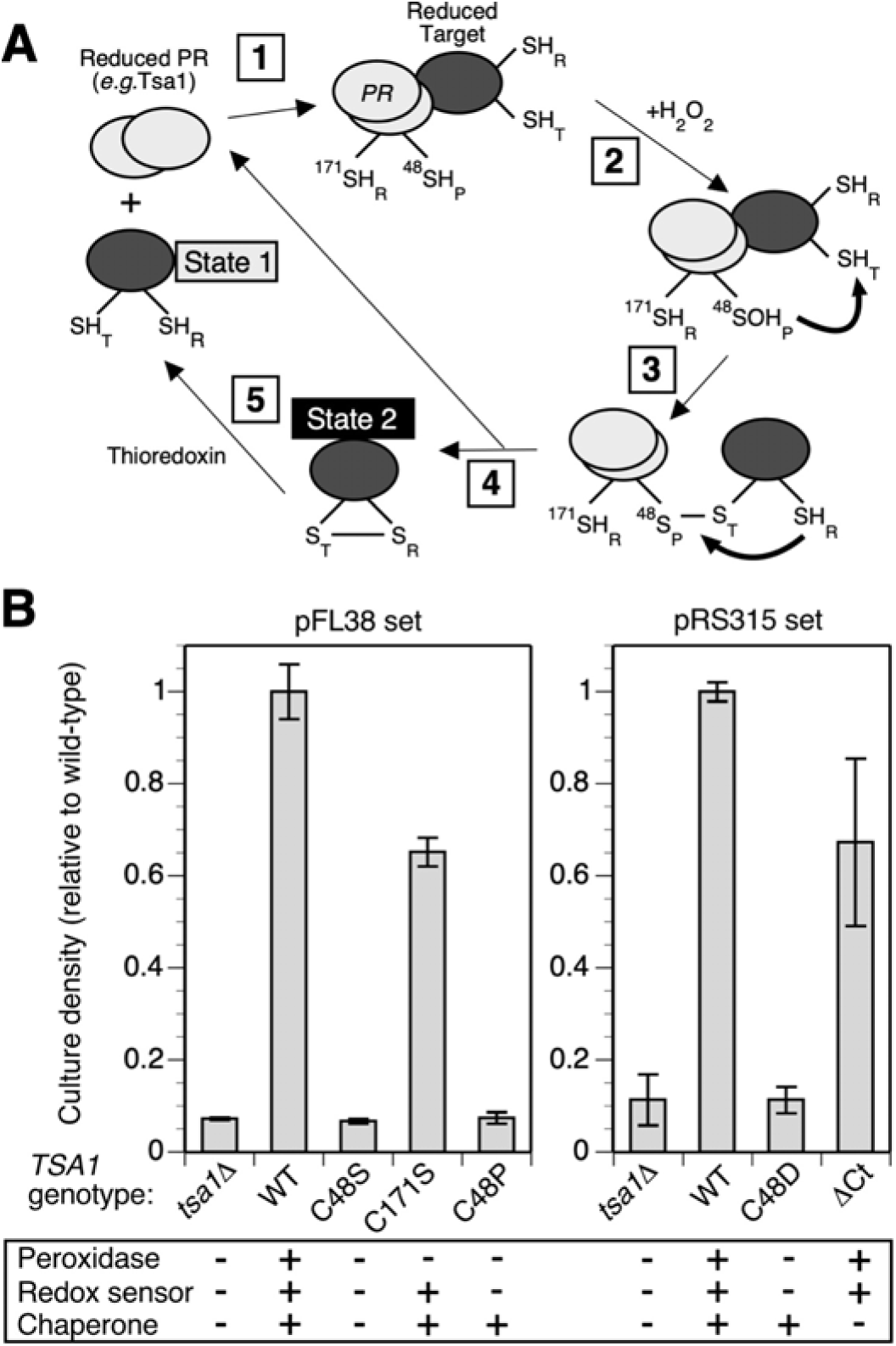
Tsa1 redox sensor activity is required for growth in low zinc. A) Scheme for PR (*grey* ovals) redox sensor function, showing PR dimer (*e.g.* Tsa1, *light grey* ovals) peroxidatic (SH_P_) and resolving (SH_R_) cysteines, and PR regulatory target (*dark grey* oval) target (SH_T_) and resolving (SH_R_) cysteines. State 1 and 2 indicate regulatory redox states of the partner (see text for details). B) Growth of strains expressing *TSA1* mutant alleles in low zinc. Table below the graphs shows the effect of the mutations on the three Tsa1 activities (peroxidase, redox sensor and chaperone). BY4742 *tsa1Δ* cells expressed the indicated wild-type or mutant forms of Tsa1 from pFL38- (*left* panel) or pRS315-based (*right* panel) vectors. Aliquots of LZM +1 μM ZnCl_2_ were inoculated to an initial *A*_595_ of 0.01 and cultures were grown for 2 days. Final *A_595_* values were normalized to the WT value for each set of strains. Data points are the average of three replicates and error bars indicate ± 1 S.D.

PR redox sensor function may provide an elegant solution to a central problem in redox regulation, how redox signals are accurately and efficiently targeted to regulated proteins [38]. The second messenger function of H_2_O_2_ is now widely recognized [57], and specific protein oxidation in response to H_2_O_2_ exposure well documented [52, 58]. However, the limited dissociation of cysteine at cytosolic pH restricts the availability of the H_2_O_2_-reactive thiolate anion, and although the intrinsic reactivity of cysteine can be increased by its molecular environment (*e.g.,* in glyceraldehyde-3-phosphate dehydrogenase) [59], relatively few such reactive cysteines have been convincingly identified. In addition, dedicated thiol peroxidase enzymes are both highly abundant and extremely reactive [37], meaning they effectively deplete ROS and RNS before such species can oxidize less reactive thiols. One model compatible with these observations is that PR’s facilitate redox regulation by transfer of oxidizing equivalents to less reactive proteins. Evidence for this view comes from various studies showing that PR’s actually promote oxidation of a subset of the proteome [38, 52, 53, 60]. One study showed that inactivating all thiol peroxidases in yeast decreased cysteine oxidation in response to peroxide treatment, and also prevented the downregulation of ribosomal gene transcription that normally occurs in response to peroxide [53]. Despite these advances however, the full scope and importance of PR redox sensor function is still unclear [38, 52].

We recently identified a mutation (*cdc19^S492A^*) that allowed a *tsa1Δ* mutant to better tolerate zinc deficiency [20]. *CDC19* encodes glycolytic pyruvate kinase (Pyk1), and *cdc19^S492A^* decreased Pyk1 activity, suggesting that Tsa1-mediated inhibition of Pyk1 contributed to adaptation. In ZnD cells, *cdc19^S492A^* increased phosphoenolpyruvate (PEP) accumulation, and alleviated the aromatic amino acid auxotrophy exhibited by a ZnD *tsa1* mutant. These data are consistent with a model in which Tsa1 downregulates Pyk1 in ZnD cells in order to spare PEP for the chorismate pathway of aromatic amino acid synthesis. Previous work showed that Tsa1 and Pyk1 physically interact *in vitro*, and that a Tsa1 C171S mutant accumulated as a mixed disulfide with Pyk1, suggesting the action of a Tsa1-Pyk1 redox relay [61]. These observations pointed to a role for the redox sensor function of Tsa1 in adaptation to low zinc. In this work, we reexamined the importance of the three Tsa1 activities for ZnD yeast, and found that the redox sensor was most critical. To identify novel targets of Tsa1 redox regulation in ZnD cells, we developed a method that effectively preserved redox-sensitive non-covalent Tsa1 interactions. Analysis of the Tsa1 interactome identified many novel targets and highlighted important processes as candidates for Tsa1 regulation, including glycolysis and amino acid synthesis. Excitingly, the Zap1 transcriptional activator was a major target for Tsa1-mediated oxidation, and Tsa1 was essential for full Zap1 activity under conditions of severe zinc deficiency. Tsa1-mediated redox regulation of Zap1 may shape the expression of the zinc regulon in response to other environmental signals.

## RESULTS

We previously reported indirect evidence for a role of Tsa1’s chaperone function in the adaptation of cells to low zinc. The Tsa1 C48S mutation inactivates both the peroxidase and chaperone activities of Tsa1, while C171S inactivates only the peroxidase activity. A Tsa1^C171S^ mutant allele complemented the growth defect of a *tsa1Δ* mutant in low Zn, suggesting that only the chaperone activity was essential [9]. However, concurrent studies on the genetic determinants of PR function [62, 63], as well as the growing appreciation for the importance of the redox sensor function of PR’s prompted us to further test this hypothesis. We determined the ability of several additional mutant alleles lacking one or more of the three known functions of Tsa1 to complement the growth defect of a *tsa1* deletion mutation in low Zn (**Fig. 1B**). Mutation of the peroxidatic cysteine (C48) to an aspartic acid mimics hyperoxidation [63], generating a chaperone-active protein that lacks peroxidase and sensor functions. Replacement of C48 with proline triggers a change in conformation with a similar effect [62]. Lastly, deletion of a conserved YF dipeptide motif at the C-terminus prevents activation of the chaperone activity by impairing hyperoxidation of C48 [64]. We confirmed that this ΔCt mutation prevented hyperoxidation by H_2_O_2_ **(Supplemental Fig. S1A**). None of the mutations markedly affected Tsa1 accumulation in either zinc-replete (ZnR) or ZnD conditions (**Supplemental Fig. S1B**).

Only the C171S and ΔCt mutants complemented the *tsa1Δ* mutation (**Fig. 1B**), indicating that only the redox sensor function is essential for ZnD cells. Notably, the chaperone-activating C48D and C48P mutants (which lack peroxidase or sensor activity) did not complement, arguing that chaperone activity alone cannot support growth in low zinc.

The importance of the Tsa1 redox sensor function suggested that Tsa1 redox-regulates partner proteins essential for growth of ZnD cells. A possible five-step model for this process is shown in **Figure 1A**. First, Tsa1 physically associates with specific reduced proteins (State 1). Second, an environmental signal triggers an increase in the level of H_2_O_2_, which reacts with the peroxidatic cysteine to form a sulfenic acid. Third, this derivative can attack a nearby cysteine on a partner protein to form a transient intermolecular “mixed” disulfide. In step 4, this linkage undergoes disulfide exchange with a resolving cysteine on the partner protein to generate an intramolecular disulfide in the partner (State 2) and modulate its function. Finally, the oxidation-dependent change in function is reversed by reduction of the partner, for example via thioredoxin activity. Although variations on this model are possible, a common prediction is that in order to maximize regulatory efficiency and specificity, Tsa1 and its partners will physically associate prior to the peroxide signal. Accordingly, we reasoned that identifying proteins non-covalently associated with Tsa1 would reveal new candidates for redox regulation.

Because previous studies of Tsa1 interactions identified relatively few partners [36, 44, 61], we suspected that the methods used might not have preserved Tsa1 interactions. We envisage two major explanations for the loss of Tsa1 interactions: First, interactions may be redox sensitive and inhibited by oxidation occurring *in vitro*; and second, use of some affinity tags might inactivate or inhibit Tsa1 binding, or require the use of complex and overlong purification procedures that dissociate weaker interactions. In response to these concerns, we developed a protocol to allow rapid and gentle isolation of Tsa1 directly from yeast cell lysates by fusing it to the maltose binding protein (MBP). MBP was reported to enhance the stability of fused proteins [65], and contains no cysteines that might react with Tsa1 or its partners. Tsa1 function can be negatively affected by modification of the C-terminus, as the addition of GFP caused both aberrant redistribution into cytosolic foci and decreased solubility [36, 66] (**Supplemental Fig. S2 B-D**). In contrast, fusing MBP to the Tsa1 C-terminus had little effect either on Tsa1 ability to support growth of ZnD cells ( **Supplemental Fig. S2A**), or its solubility (**Supplemental Fig. S2 B-D)**. Finally, to inhibit the oxidation of Tsa1 or its partners *in vitro* and preserve redox-sensitive interactions, yeast cells expressing Tsa1-MBP were pretreated with the thiol-reactive alkylating agent N-ethylmaleimide (NEM) prior to lysis, and NEM was routinely included during subsequent purification steps (see Experimental Procedures).

A previous study used the C171S mutant of Tsa1 to capture potential regulatory partners [61]. The rationale for this approach was that mutation of the resolving cysteine blocked its reaction with sulfenylated Tsa1C48 and thus promoted the alternative formation of mixed disulfides with associated proteins. For this reason, we also used the Tsa1^C171S^ allele fused to MBP in our initial fractionation experiments (abbreviated TRM, for Tsa1 Resolving cysteine Mutant). Proteins specifically associated with TRM were identified by comparison with cells expressing MBP alone. Immunoblotting of crude protein extracts showed that MBP and TRM were expressed at similar levels in ZnR and ZnD cells (**Fig. 2A, B**). After purification of MBP or TRM, co-purifying proteins in eluate fractions were visualized by silver staining (**Fig. 2B**). MBP-alone samples contained a prominent band at the expected size for MBP, along with minor amounts of other proteins. In contrast, TRM fractions contained an 82 kDa band corresponding to TRM and many proteins not observed in the MBP fraction. Detection of MBP and TRM in immunoblots of eluate fractions (**Fig. 2C**) showed that TRM fractions contained slower-migrating reduction-sensitive forms, which likely represent mixed disulfides of TRM and other proteins. Because the cells were treated with NEM both prior to and during protein extraction and purification, these redox-sensitive forms were likely present *in vivo*. Indeed, subsequent experiments confirmed that mixed disulfides of Tsa1^C171S^ were also present in cell extracts prepared under conditions that fully preserved redox state (*e.g.,* see **Fig. 10)**.

**Figure 2.**
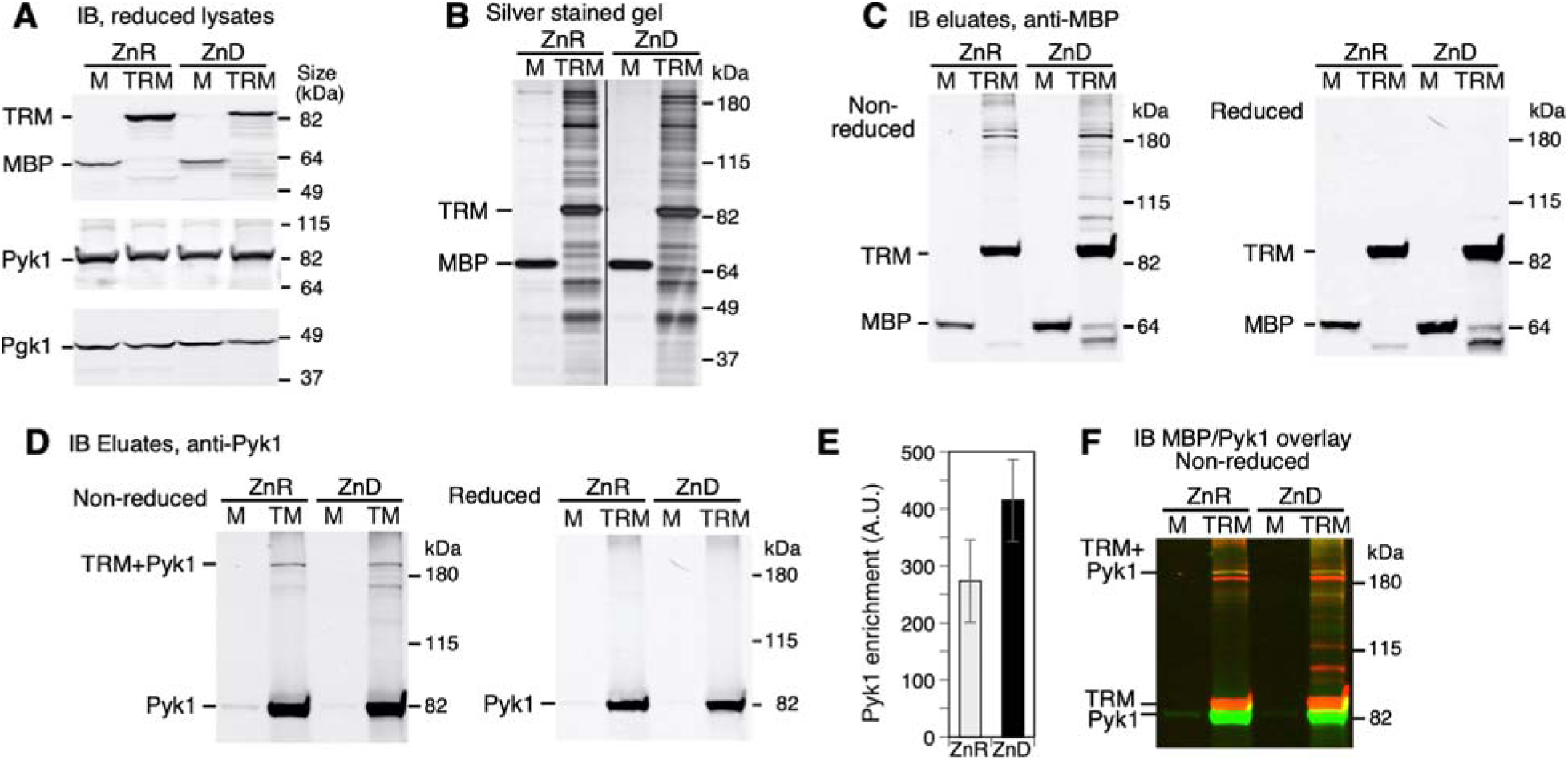
Tsa1^C171S^-MBP associated with a mixture of proteins including Pyk1. A) Wild-type yeast (BY4742) expressing MBP alone (M) or congenic *tsa1Δ* mutant expressing Tsa1^C171S^-MBP (TRM) were grown to log phase in LZM + 100 μM ZnCl_2_ (ZnR) or LZM + 1 μM ZnCl_2_ (ZnD). Protein was extracted and equal amounts were subjected to SDS-PAGE and immunoblot analysis to detect MBP, Pyk1, and Pgk1. Note that in this and subsequent immunoblot figures, bands detected with anti-MBP in TRM lanes that are lower than 82 kDa molecular mass (*i.e.* the size of the fusion protein) are due to minor degradation during protein preparation. B) MBP or TRM were purified from equal amounts of lysate protein isolated from the strains described in (A), and equal volumes of column eluate were separated by SDS-PAGE and visualized by silver staining. Vertical lines in this and the subsequent panels and immunoblot figures indicate removal of extraneous lanes. C, D) Eluate samples shown in (B) were reduced with DTT (*right* panels) or left untreated (*left* panels) and subjected to immunoblot analysis with antibody to MBP (C) or Pyk1 (D). E) Pyk1 enrichment in reduced TRM fractions relative to MBP alone. Values are the average of three replicates and error bars indicate ± 1 S.D. F) Overlay of non-reduced MBP (C) and Pyk1 (D) immunoblots, indicating the disulfide crosslinked complex of TRM and Pyk1 (*yellow* band > 180 kDa). For all gels and IB, size of molecular mass standards are indicated on right.

### Pyk1 physically associates with TRM

To further validate the function and utility of TRM, we characterized its interaction with the glycolytic enzyme pyruvate kinase isozyme 1 (Pyk1), a suspected target of Tsa1 regulation [20, 61]. A previous study showed that a fraction of a Tsa1^C171T^ resolving cysteine mutant was disulfide crosslinked to Pyk1 [61] and that Tsa1 and Pyk1 physically interacted *in vitro*. Accordingly, we found that while cells expressing TRM or MBP alone accumulated similar amounts of Pyk1 (**Fig. 2A)**, Pyk1 was enriched at least 250-fold in purified TRM eluate fractions vs MBP alone (**Fig. 2D, E**). Immunoblotting of non-reduced TRM eluate fractions revealed that Pyk1 co-purified as two major species, an abundant band of 80 kDa apparent molecular weight, and a lower abundance species at >180 kDa (**Fig. 2C, D, F**). The 180 kDa species was reduction sensitive and reacted with both antibodies, indicating it represented a mixed disulfide of Pyk1 and Tsa1^C171S^ **(Fig. 2F**, >180 kDa *yellow* band), likely equivalent to the species previously detected by immunoprecipitation of myc-tagged Tsa1 [61]. In contrast to this report however, we found that the great majority of Pyk1 that co-purified with Tsa1^C171S^ was not crosslinked to it (94.9 +/- 2.6% S.D., **Fig. 2D**).

### Pyk1-Tsa1 association does not depend on Tsa1 chaperone activity

The prominent non-covalent interaction of Tsa1^C171S^ and Pyk1 prompted us to examine the functional requirements for this association. Tsa1 has three major functions: a thioredoxin-dependent peroxidase (which requires both Tsa1 C48 and C171 thiols), a redox sensor (which requires only the C48 thiol), and a holdase-type protein chaperone (which is activated by C48 hyperoxidation and thus requires C48 but not C171). We initially supposed that preparation of cell extracts under aerobic conditions generated excess ROS that hyperoxidized Tsa1 (**Fig. 3A)**, and that the resulting increase in Tsa1 chaperone activity promoted its non-covalent interaction with Pyk1. However, even when native cell lysates were prepared without NEM treatment, we detected no hyperoxidation of wild-type Tsa1-MBP by immunoblotting (**Fig. 3B**, reduced lysates, lane 5). Hyperoxidation was only detected when cells were pretreated with a high concentration of H_2_O_2_ prior to lysis (**Fig. 3B**, reduced lysates, lane 8). The antibody used to detect hyperoxidized Tsa1 was specific, as it did not detect the Tsa1^C48S^ mutant in H_2_O_2_-treated cells (**Fig. 3B**, reduced lysates, lane 9). We then examined the effect of enhancing Tsa1 hyperoxidation on Pyk1 association. H_2_O_2_ treatment efficiently induced hyperoxidation, as indicated by the near absence of oxidatively crosslinked Tsa1-MBP dimers (**Fig. 3A, B**, non-reduced eluates, compare lanes 5 and 8). However, hyperoxidation did not promote the association of Tsa1-MBP and Pyk1 (**Fig. 3B**, reduced eluates with α-Pyk1, compare lanes 2 and 8). To further examine the role of the chaperone function in the Pyk1-Tsa1 interaction, we compared Pyk1 association with MBP-tagged Tsa1 mutants lacking specific Tsa1 activities (C48S, C171S, and ΔCt) **(Fig. 3C, D**). Both the C48S and ΔCt mutations prevent chaperone activation by preventing C48 hyperoxidation (**Fig. 3A, Supplemental Fig. S1A**). These mutations either increased Pyk1 association (C48S, **Fig. 3C**) or had no effect (ΔCt, **Fig. 3D**), indicating that chaperone activity was not required for Pyk1 association. Together, these observations further indicate that the prominent non-covalent interaction of Tsa1 and Pyk1 was independent of chaperone activity.

**Figure 3.**
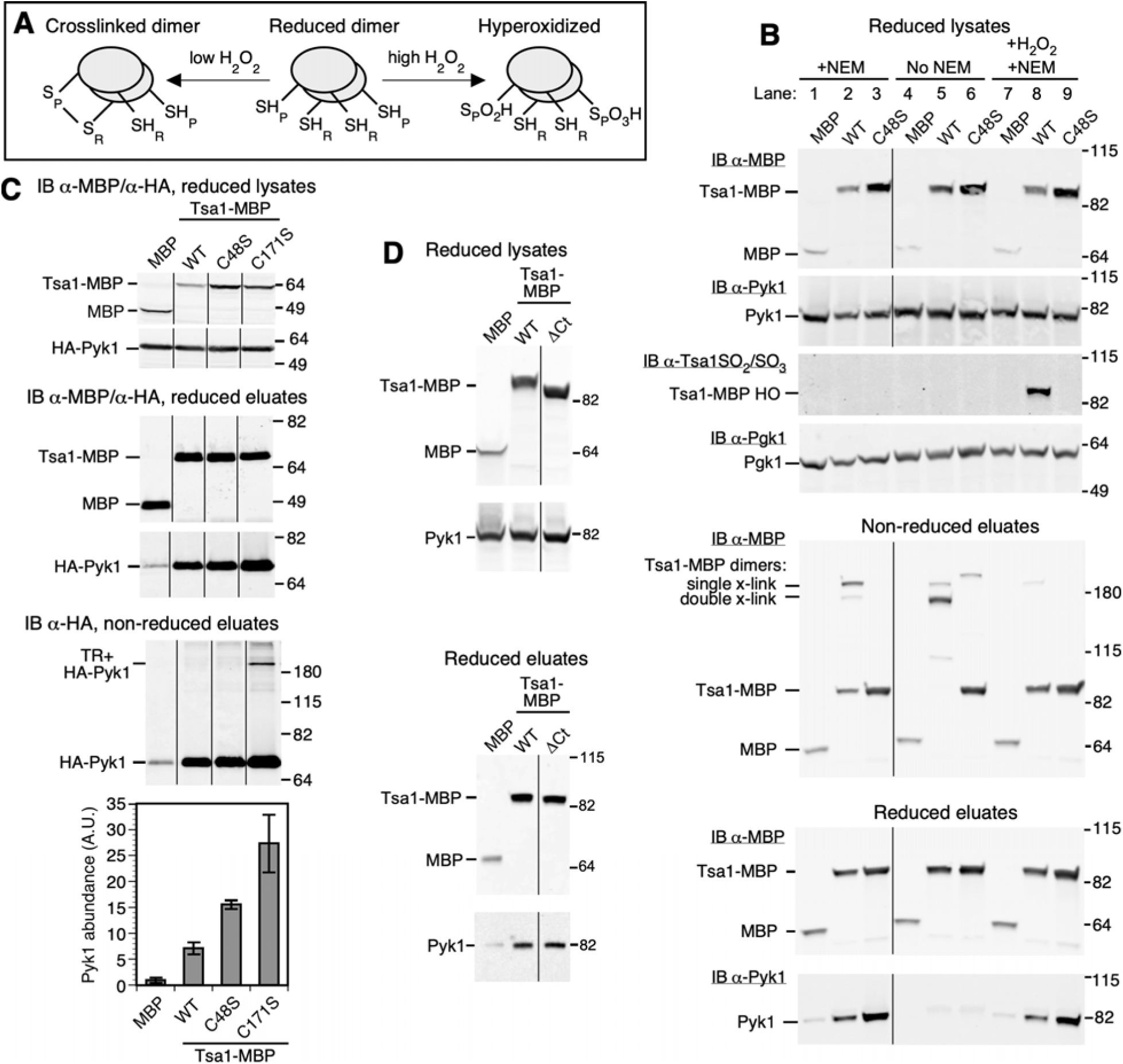
Pyk1 and Tsa1 association is independent of Tsa1 chaperone activity. A) Effect of substrate on Tsa1 oxidation state. At low substrate concentration (*left*), the peroxidatic cysteine (SH_P_) of Tsa1 is oxidized to a sulfenic acid, which can then react with the resolving cysteine of the partner monomer (SH_R_) to form single or double disulfide-linked dimers. Excess substrate (*right*) prevents crosslinking by driving SH_P_ hyperoxidation to a sulfinic or sulfonic acid, activating chaperone activity. B) Effect of NEM on the non-covalent association of Pyk1 with Tsa1-MBP. Wild-type yeast (BY4742) expressing MBP alone (M) or congenic *tsa1Δ* mutant expressing Tsa1-MBP (WT) or Tsa1^C48S^-MBP (C48S) from plasmids were grown to log phase in LZM+100 μM zinc. Cell lysate and purified MBP fractions (amylose column eluates) were subject to immunoblotting to detect MBP, Pyk1, and Pgk1 (as loading control) with specific antibodies as indicated. Hyperoxidized Tsa1 (Tsa1-HO) was detected with an antibody to sulfinylated or sulfonylated Tsa1. Where indicated (+NEM), cells were treated with 10 mM NEM prior to cell lysis, and all purification buffers contained 10 mM NEM. “No NEM” indicates cells were not NEM treated and NEM was omitted from buffers. Some cells were also treated with 1 mM H_2_O_2_ for 5 min (+H_2_O_2_), prior to NEM treatment of cells. Displayed results are representative of three independent experiments. C) Effect of Tsa1 C48S and C171S mutations on Pyk1 association. Wild-type yeast (BY4742) expressing MBP alone (M) or congenic *tsa1Δ* mutant expressing Tsa1-MBP (WT), Tsa1^C48S^-MBP (C48S), or Tsa1^C171S^-MBP (C171S) from plasmids were grown to log phase in LZM+100 μM zinc. Extracts of cells were prepared in buffer supplemented with 5 mM NEM, MBP was purified, and MBP-tagged proteins or Pyk1 were detected as described for panel B. Graph shows recovery of reduced Pyk1 quantified from three experimental replicates and the error bars indicate ± 1 S.D. Note use of 10% acrylamide gels for these immunoblots resulted in differences in apparent protein molecular mass vs. the marker proteins when compared with 4-15% gradient gels (panels B and D). Positions of the marker bands are indicated for all immunoblots. D) Effect of Tsa1 ΔCt mutation on Pyk1 association. Zinc-replete cultures of wild-type cells (BY4742) expressing MBP alone, or congenic *tsa1Δ* cells expressing wild-type Tsa1-MBP (WT) or Tsa1^ΔCt^-MBP (ΔCt) from plasmids were treated with 10 mM NEM prior to protein extraction (MBP purification buffers also contained 10 mM NEM). MBP and Pyk1 were detected in immunoblots of reduced lysates (*top* panel) and reduced MBP fractions (eluates, *lower* panel). Displayed results represent two independent experiments.

During these experiments we observed that in the absence of NEM treatment, the Pyk1-Tsa1 non-covalent association strongly decreased (**Fig. 3B**, reduced eluates, lanes 2 and 3 vs. 5 and 6). Under these conditions Tsa1 was efficiently oxidized, as cell lysates prepared aerobically predominantly contained Tsa1 disulfide-linked dimers (**Supplemental Fig. S3A)**, while samples prepared using TCA to prevent oxidation primarily contained monomeric Tsa1 **(Supplemental Fig. S3B)** (note that oxidized Tsa1 dimers can be crosslinked once or twice via C48-C171 linkages, and these two forms migrate differently in SDS-PAGE). NEM treatment prior to and during cell fractionation inhibited Tsa1 dimer formation (**Fig. 3B**, non-reduced eluates), and also preserved Tsa1-Pyk1 association (**Fig. 3B**, reduced eluates), suggesting Tsa1 oxidation *in vitro* triggered its dissociation from Pyk1. Since the hyperoxidation of C48 *in vivo* did not inhibit this interaction (**Fig. 3B)**, we suggest that Tsa1 peroxidase cycling is responsible for dissociating the interaction, perhaps because this cycle requires that Tsa1 associate with thioredoxin for its reduction. Consistent with this interpretation, mutations that eliminated Tsa1 peroxidase cycling (C48S and C171S) substantially increased Pyk1 association (**Fig. 3C**). However, as NEM treatment also enhanced Pyk1 association with these Tsa1 mutants (**Supplemental Fig. S3C)** and Pyk1 oxidation also occurs during cell lysis [20], these observations also suggest that the redox state of Pyk1 influences its interaction with Tsa1. This apparent preference of Tsa1 for reduced Pyk1 is an interesting feature which, if it extends to other Tsa1-target interactions, might be expected to enhance the overall efficiency of redox regulation.

*Tsa1^C171S^ associates with Pyk1* in vivo. It was conceivable that the formation of disulfide-linked and/or non-covalent complexes of Pyk1 and Tsa1 occurred during cell lysis and purification, rather than *in vivo*. To address this issue, we used the cell-permeant crosslinking agent formaldehyde (FA) to examine Tsa1 and Pyk1 association *in vivo*. Cells expressing HA-tagged Pyk1 and either MBP or TRM were treated with 1% FA for 10 min to minimize non-specific crosslinking [67]. FA was then quenched with glycine, protein extracted, and MBP or TRM purified. Proteins were reduced with DTT and/or heated to reverse FA crosslinks before immunoblotting with anti-MBP and anti-HA antibodies. Immunoblots of non-reduced TRM eluate samples from non-FA treated cells (**Fig. 4A**, *left* panel, lane 3) showed multiple bands with apparent molecular weight higher than TRM. These were DTT-sensitive indicating they were potential mixed disulfides (**Fig. 4B**, *left* panel, lane 3). FA treatment of cells increased the number and abundance of high molecular weight TRM species (**Fig. 4A**, *left* panel, lane 4 vs. 3). Reduction of these FA-treated samples (**Fig. 4B**, *left* panel, lane 4) revealed a residual collection of FA-crosslinked species, as indicated by their sensitivity to heat treatment (**Fig. 4B**, *left* panel, lane 8). These species likely represent FA-crosslinked multimers of TRM as well as TRM crosslinked to other proteins *in vivo*.

**Figure 4.**
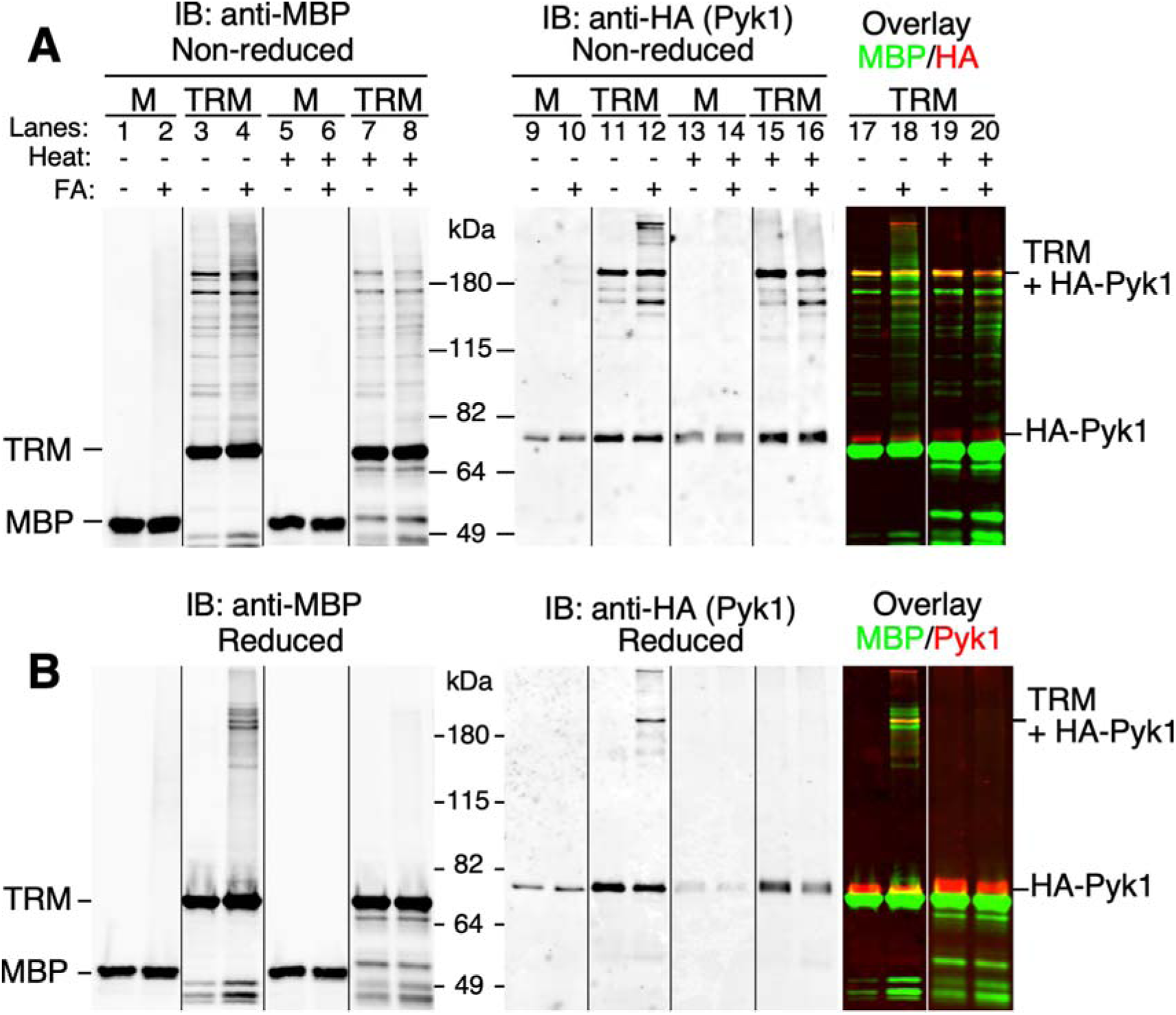
Tsa1 and Pyk1 associate *in vivo*. Wild-type (BY4742) yeast cells expressing MBP, or congenic *tsa1Δ* cells expressing Tsa1^C171S^-MBP (TRM) were grown to log phase in ZnD media and treated with 1% formaldehyde (FA) for 10 min (a duplicate set of samples not FA treated are also shown). FA was quenched with glycine, cell lysates prepared, and MBP purified. To promote Tsa1-Pyk1 mixed disulfide formation *in vitro,* pretreatment of cells with NEM was omitted, but 5 mM NEM was included in fractionation buffers. Prior to immunoblotting, aliquots of some samples were reduced by addition of SDS-PAGE sample buffer (SB) with 10 mM DTT followed by incubation at 37° for 1 h (B). To reverse FA crosslinks alone, SB with no DTT was added and samples heated to 95° for 10 min (Heat). To reverse both types of crosslinks, SB+DTT was added and samples heated to 95° for 10 min. Samples were fractionated on 10% gels before immunoblotting. Note bands below TRM are minor degradation products.

Detection of HA-tagged Pyk1 in TRM eluate samples (**Fig. 4A**, *right* panel) revealed a Pyk1 band of approximately 70 kDa, as well as several species migrating above the 115 kDa marker. One of these was the previously observed mixed disulfide with TRM, as it was also detected with the MBP antibody (TM+HA-Pyk1, *yellow* band >180 kDa in overlay). FA treatment of cells also generated additional high molecular weight species of Pyk1, which were eliminated by heat treatment (**Fig. 4A**, compare lanes 12 and 16). Heat treatment alone had little effect on the TM+HA-Pyk1 mixed disulfide, but this band was completely lost when samples were both reduced and heat treated **(Fig. 4B**, lane 16), indicating it represented Pyk1 that was both disulfide and FA crosslinked to TRM. These experiments thus demonstrate that Pyk1 and Tsa1^C171S^ physically associate *in vivo*. Further evidence for an *in vivo* association came from experiments with wild-type Tsa1-MBP (**Supplemental Fig. S4**). When cells were exposed to a low concentration of H_2_O_2_ (0.1 mM), then treated with NEM to halt oxidation, a redox-sensitive form of Pyk1 was generated with an apparent size identical to the mixed disulfide formed with TRM. Formation of this species is an early step in a redox relay between wild-type Tsa1 and Pyk1 (**Fig. 1A**) that requires physical association of these factors.

### Tsa1 specifically oxidizes C174 of Pyk1

If Tsa1 provides a “redox sensor” function for Pyk1, we expected that Tsa1 would specifically react with a single Pyk1 cysteine (the “target”) (**Fig. 1A**), directly or indirectly responsible for modulating Pyk1 activity. To test this prediction, we mutated each of the seven Pyk1 cysteine residues to alanine, and determined the effect on formation of the mixed disulfide with TRM. Only Cys174 was essential for the accumulation of this species (**Fig. 5A**). Cys174 is a surface exposed residue located in the Pyk1 B-domain, a flexible region that acts as a “lid”, closing over the A-domain active site when substrate is bound (**Fig. 5B**) [68]. The B-domain is an essential component of the active site that directly contributes to binding the substrate ADP via a conserved arginine residue (Arg91 in Pyk1) [68]. These observations suggest that oxidation of C174 might inhibit Pyk1 function by restricting the movement of the B-domain. We note that this effect could result from formation of the mixed disulfide itself, by resolution of this complex to form an intramolecular disulfide in Pyk1, or by crosslinking of Pyk1 subunits within a multimeric complex.

**Figure 5.**
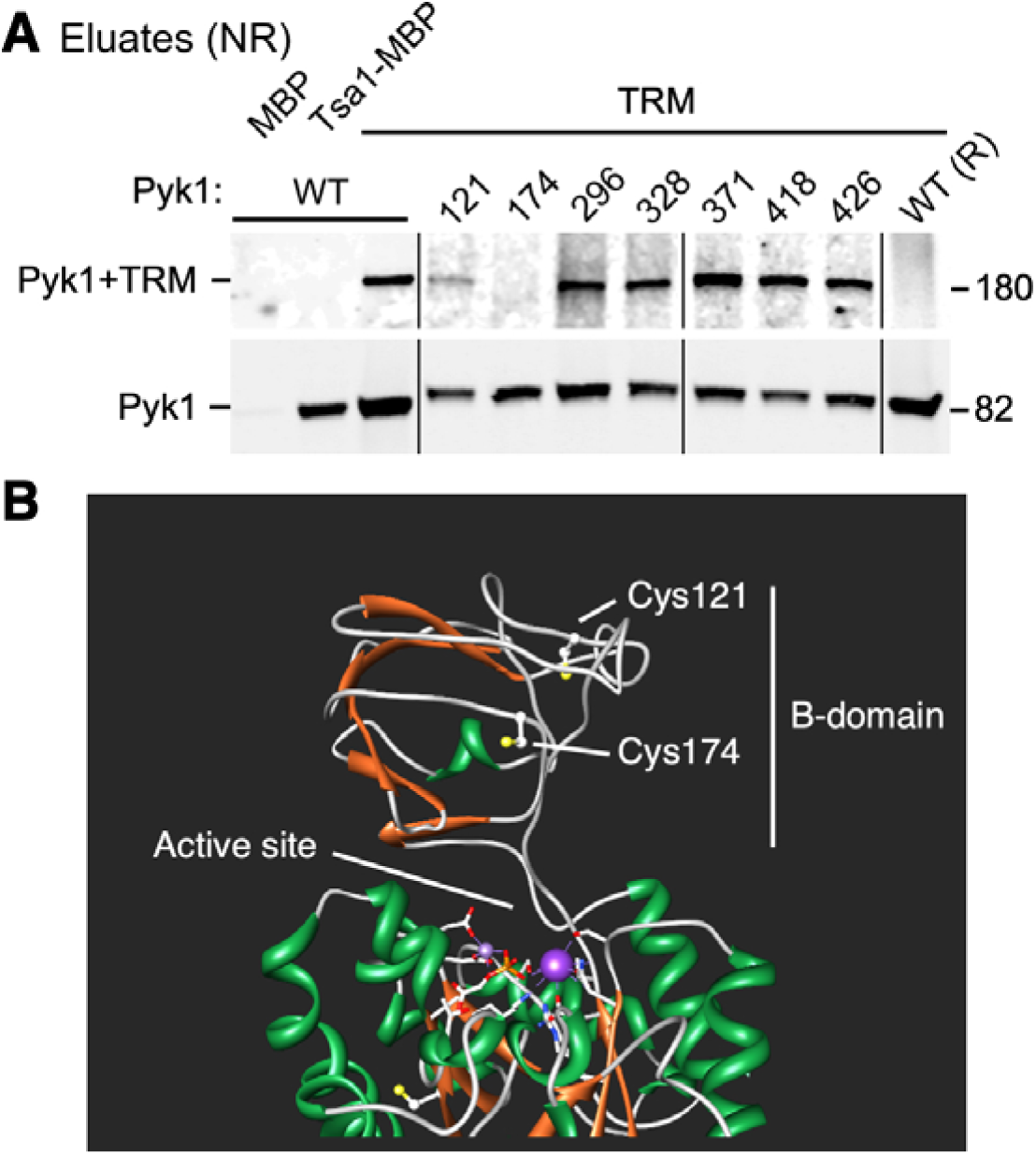
Oxidative crosslinking of Pyk1 and TRM is dependent on Pyk1 cysteine 174. A) A set of *tsa1Δ* mutant strains carrying each of seven cysteine to serine mutations at the *CDC19* (Pyk1) genomic locus (CWM357, 359, 361, 363, 365, 367, and 442, **Supplemental Table S8**) were transformed with the pTsa1^C171S^-MBP plasmid (TRM). In addition, a *CDC19* wild-type strain (BY4742) was transformed with pMBP, and a congenic *tsa1*Δ mutant was transformed with wild-type pTSA1-MBP or TRM as indicated. Strains were grown in ZnR medium, treated with 10 mM NEM, and protein lysates prepared for MBP purification. Non-reduced eluate samples were subject to immunoblotting to detect Pyk1 crosslinked to TRM (*upper* panel) and uncrosslinked Pyk1 (*lower* panel). A duplicate *CDC19 tsa1Δ*/TRM sample (see lane 3) was reduced with DTT prior to electrophoresis (R, last lane). Lines indicate where superfluous lanes were removed. B) Cysteine 174 is located in the B-domain near the Pyk1 active site. A single subunit of Pyk1 (PDB model 1A3X) is shown indicating the positions of the two cysteine side chains in the B domain relative to the active site with bound substrate.

In summary, these data indicated that Tsa1 physically interacted with Pyk1 *in vivo*, that this novel non-covalent interaction was sensitive to Tsa1 and Pyk1 oxidation, and that it was not dependent on Tsa1 chaperone activity. Additionally, we found that wild-type Tsa1 forms oxidatively crosslinked species with Pyk1 in response to H_2_O_2_, suggesting this redox relay is physiologically relevant, and determined that cysteine 174 of Pyk1 is essential for its oxidation by Tsa1. Together, these observations provided further support for the involvement of Tsa1 in a redox relay for Pyk1 regulation. More generally, they indicated that TRM is a useful reagent for the identification of novel Tsa1-interacting factors.

### Identification of the Tsa1 interactome

Our observation that the C171S mutation enhanced the non-covalent interaction of TRM and Pyk1 suggested that an analysis of the Tsa1^C171S^ interactome would identify novel Tsa1-associated proteins and potential regulatory targets, including those facilitating adaptation to ZnD conditions. To identify the interactome, we performed a label-free mass spectrometric comparison of MBP and TRM eluates. Cultures of wild-type cells expressing MBP, and *tsa1Δ* cells expressing TRM were grown to log phase in ZnR and ZnD conditions, protein extracted, and MBP purified. Because NEM treatment of cells was found to reduce MS detection sensitivity (data not shown), we suppressed oxidation during and after cell lysis by using the MS-compatible alkylating reagent iodoacetamide (IAA) in protein extraction and purification buffers. Under these conditions, SDS-PAGE confirmed that multiple proteins co-fractionated with TRM but not with MBP alone, and immunoblotting confirmed Pyk1 co-purification with Tsa1^C171S^ -MBP (**Supplemental Fig. S5A-D**). Treatment with IAA effectively preserved oxidation-sensitive interactions, as the distribution of protein species seen in SDS-PAGE gels of IAA-treated and purified TRM was substantially unaffected by reduction (**Supplemental Fig. S5E, F**), indicating that co-fractionating proteins were primarily non-covalently associated. To identify these proteins, column eluates of MBP and TRM samples were subjected to label-free LC-MS/MS analysis. Protein and peptide detection thresholds were set at 1% FDR with a minimum of 2 peptides per protein required for identification. Proteins were quantified from spectral counts and signal normalized using EmPAI [69], identifying many new Tsa1-interacting proteins (**Supplemental Table S1**).

To rank potential Tsa1 partners, we first calculated enrichment of proteins in ZnR TRM samples vs MBP alone. For proteins detected in TRM but not in MBP alone samples, fold enrichment was calculated by setting the minimum signal to 0.001. Excluding transposon-encoded proteins, 112 proteins were enriched at least 3-fold in ZnR TRM fractions (Students t-test, no multiple comparison correction, *p*<0.05, **Supplemental Table S2**, panel 1). This value includes 12 sets of paralogous proteins sharing identical peptides. In ZnD samples, we identified 120 proteins enriched >3-fold (**Supplemental Table S2**, panel 1). Analysis of the overlap of ZnR and ZnD data revealed 56 interacting proteins found under both conditions, another 56 unique to ZnR, and 64 unique to ZnD, for a total of 176 proteins enriched >3-fold in ZnR and/or ZnD samples. To determine how many of these targets were novel, we examined the set of Tsa1 physical interactions compiled in BioGRID [70], which includes 88 proteins reported to interact with Tsa1 **(Supplemental Table S3)**. Of these proteins, only 16 were also found in the TRM interactome, meaning 160 new candidates were identified. Twelve of the 16 previously reported interactions have strong independent experimental support **(Supplemental Table S3)**, indicating that genuine Tsa1 interactions were identified. Notably, the 62 proteins listed in BioGRID but absent from the interactome were detected in large scale studies and lack independent verification, suggesting they may represent false positives or artifacts.

### Tsa1 binding proteins identify potentially regulated processes

As an objective test of functional enrichment within the interactome, we used FunSpec [71] to examine the list of 176 significantly enriched proteins for disproportionate representation of Gene Ontology (GO) process, function, and component terms, and Munich Information Center for Protein Sequences (MIPS) functional classifications (see Experimental Procedures). More informative terms identified for each general class are listed in **Table 1** (the full analysis is provided in **Supplemental Table S4)**. This analysis revealed that multiple processes were significantly overrepresented in the Tsa1 interactome. Some classes were expected given known Tsa1 functions: *e.g.,* we identified proteins involved in redox homeostasis, several of which (*e.g.,* Tsa2, Srx1, and Trx1) were already known to interact with Tsa1. Even within this group however, several novel partners were identified (Ccs1, Sod1, Mxr1, Ahp1, and Rck2). Most are antioxidant enzymes, but Rck2 is a kinase that participates in the oxidative stress response, suggesting another mechanism by which Tsa1 could contribute to this process.

**Table 1.**
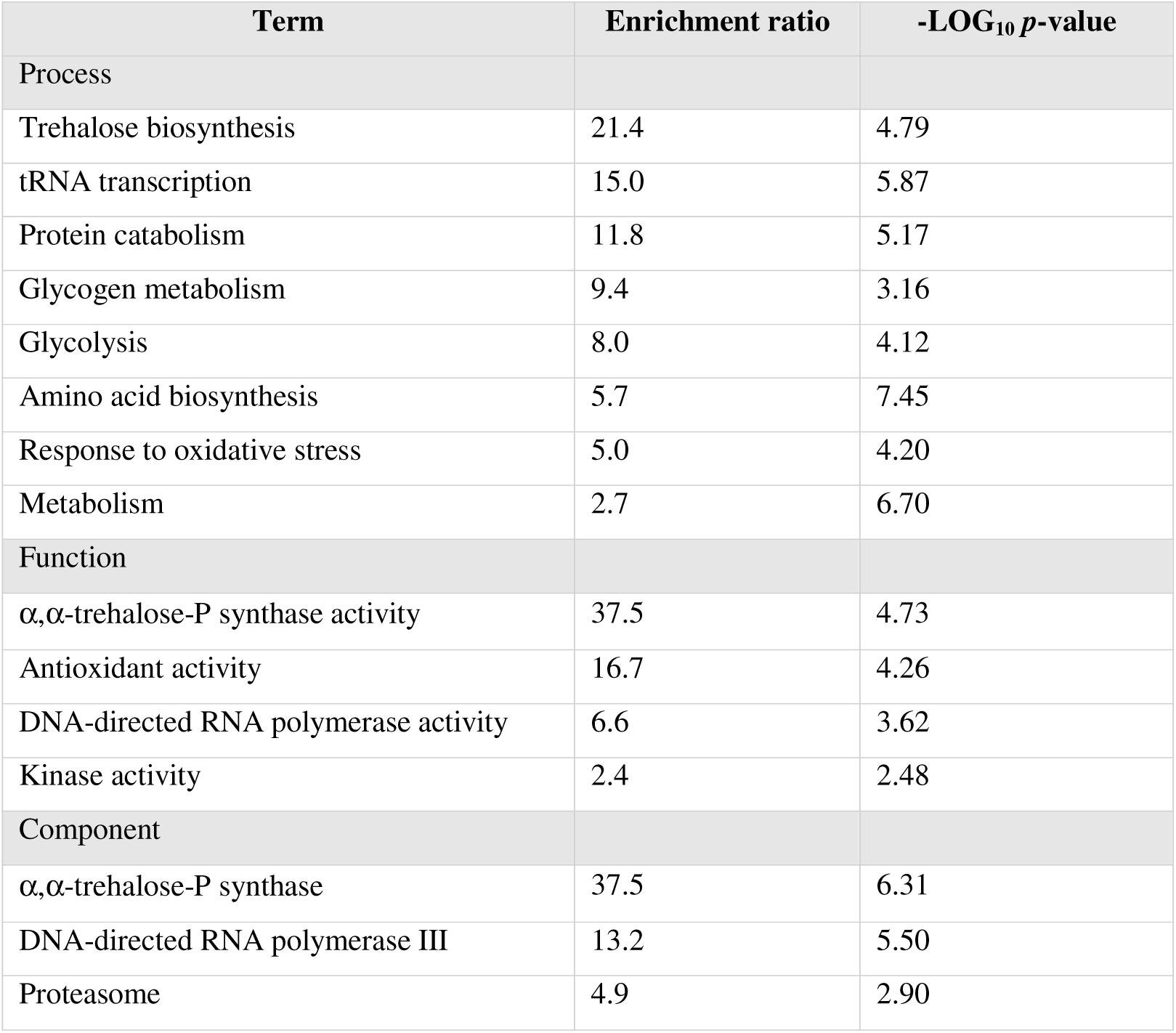
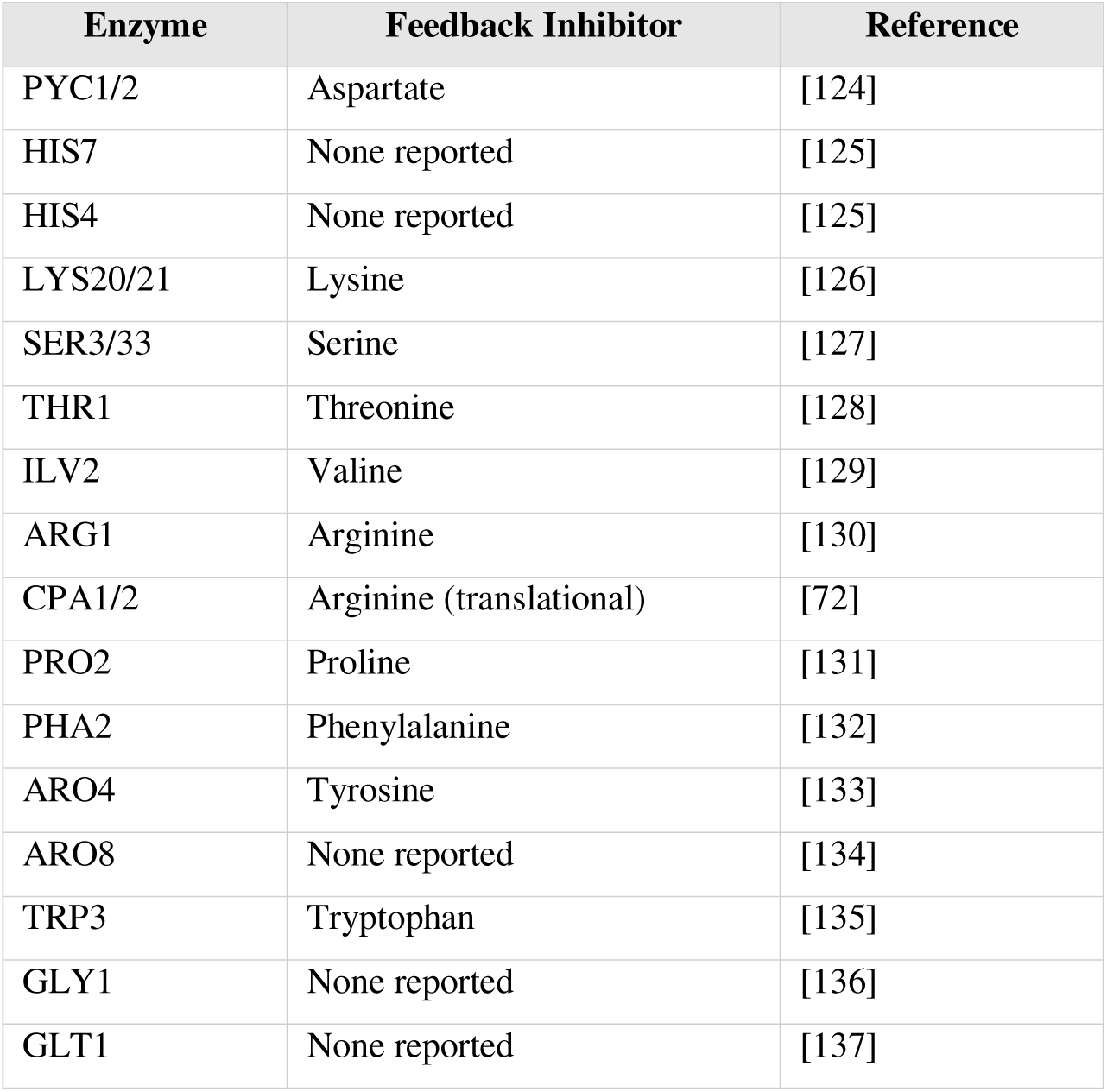
**A.** Selected GO component, function, and process terms significantly overrepresented in the Tsa1 interactome vs. the complete proteome. GO term enrichment *p*-value was calculated using Funspec. **B.** Interactome components involved in amino acid biosynthesis, indicating those feedback inhibited by pathway products.

In addition to its previously known roles, GO analysis also linked Tsa1 to a number of novel processes and pathways, including the proteasome and eisosome complexes, RNA polymerase III, protein transport via the cytoskeleton, and the mitotic cell cycle. Most notable however were GO terms related to intermediary metabolism. Components of core metabolic pathways such as glycolysis, carbohydrate synthesis, and amino acid, purine, and pyrimidine synthesis were abundant in the interactome. In glycolysis, we identified both phosphofructokinase isozymes (Pfk1 and 2) and FBP-aldolase (Fba1) as well as Pyk1. We also identified enzymes for synthesis of storage carbohydrates glycogen (Gsy1, Glg1), and trehalose (Tsl1 and Tps1-3), and enzymes representing multiple glycolysis-dependent pathways of carbohydrate metabolism, such as synthesis of glycerol (Gpd2), inositol (Inm2), and the cell wall polysaccharide chitin (Gfa1, Pcm1). Interestingly these proteins disproportionately represent highly regulated and/or committed steps in their respective pathways, which is consistent with Tsa1 providing additional regulatory input. This trend was particularly evident for the amino acid synthesis pathways, which were highly represented in the interactome (**Table 1**, **Fig. 6**, and **Supplemental Figure S6**). For the subset of these pathways that included Tsa1 targets, we observed a significant overrepresentation of the enzymes most critical to the rapid inhibition of pathways by their products (*i.e.* the enzymes allosterically inhibited by amino acids) (*p*=0.02, Pearsons Chi^2^, **Supplemental Table 5)**. Although necessarily simplified, this analysis was conservative, as proteins with multiple activities (*e.g.,* His4) were counted as a single enzyme, and other mechanisms of specific feedback inhibition were excluded (*e.g.,* the translational downregulation of Cpa1/2 by arginine, **Supplemental Figure S6**) [72]. Notably, three of the four enzymes of the chorismate pathway identified as Tsa1 targets (Aro4, Pha2 and the Trp2/3 complex) are feedback inhibited by pathway products (**Fig. 6**). In addition to conserving PEP [20], Tsa1 might also support aromatic amino acid synthesis in ZnD cells by regulation of these targets, perhaps to compensate for reduced activity of the zinc-dependent enzyme 3-DHQ synthase (Aro1).

**Figure 6.**
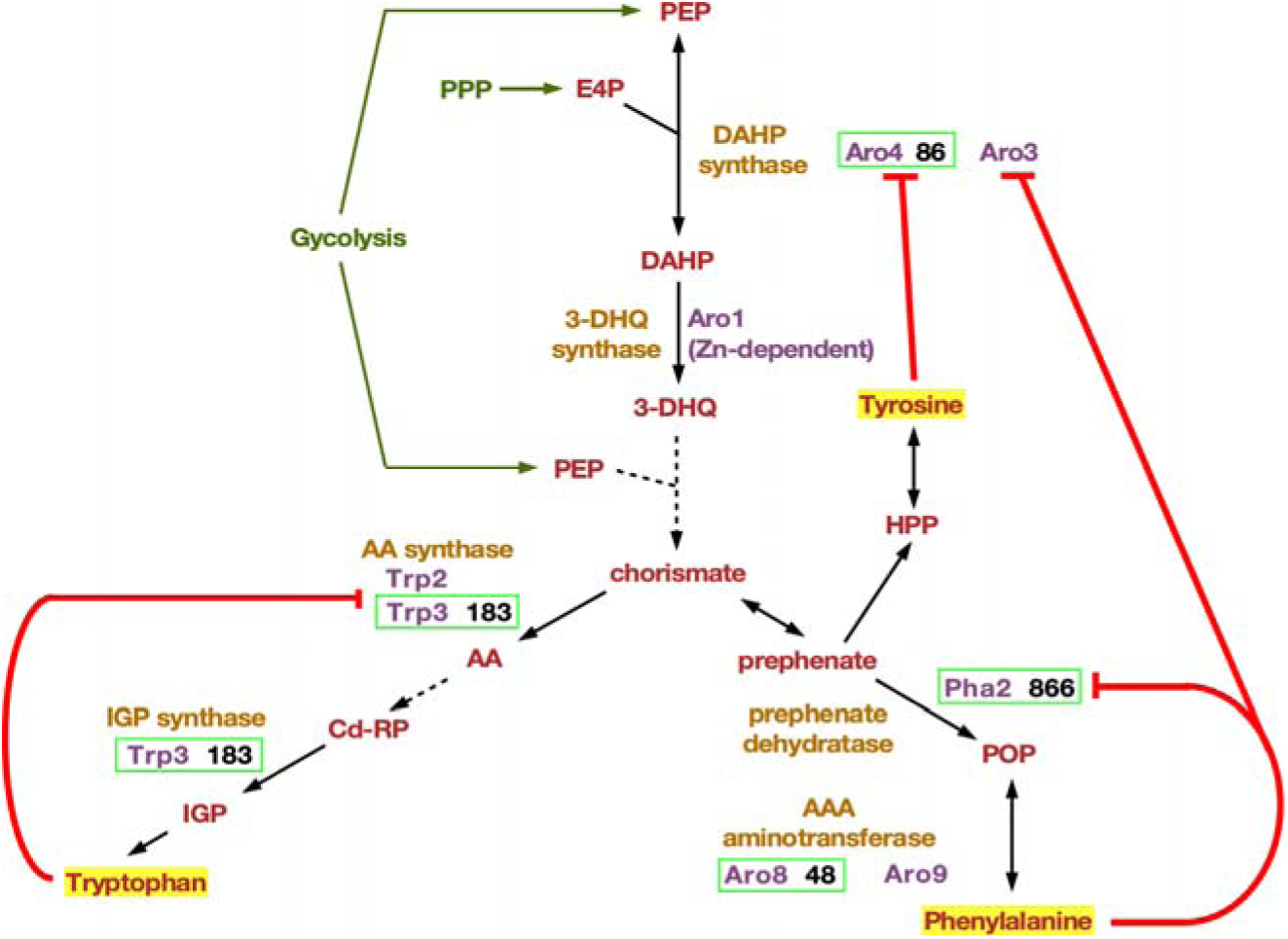
Tsa1 interactome components in aromatic amino acid pathways. Schematic of the yeast chorismate pathway showing enzymes (*purple*) and their activities (*orange*). Interactome components are outlined in *green*. Numbers indicate fold enrichment in eluates of TRM vs MBP alone from ZnD cells (see **Supplemental Table S2**). *Dashed* lines indicate multiple enzyme catalyzed transformations. *Red* lines indicate enzyme inhibition by pathway products. Abbreviations are: PEP, phosphoenol pyruvate; E4P, erythrose 4-phosphate; DAHP, 3-deoxy-arabino-heptulosonate 7-phosphate; 3-DHQ, 3-Dehydroquinate; AA, Anthranilate; CdRP, 1-(o-carboxyphenylamino)-1’-deoxyribulose 5’-phosphate; IGP, indole-3-glycerol-phosphate; HPP, 3- (4-hydroxyphenyl)pyruvate; POP, 3-phenyl-2-oxopropanoate.

### Verification of Tsa1 partner association

To validate the interactome data, a selection of potential partners from the combined ZnR and ZnD set were verified by performing independent purification and immunoblotting experiments. To minimize potential target dissociation due to oxidation, cells were treated with NEM prior to protein extraction and NEM was included in the fractionation buffer. We selected several targets for which antibodies were available, including Fba1, Pyc1/2, Pfk1/2, Hsp42, and Ccs1, and also utilized epitope- or GFP-tagged versions of Zap1, Nce103, Adh4, and Aro4. All these candidate targets were enriched in TRM eluates vs MBP alone (**Fig. 7A, Supplemental Table S2,** panel 2). In contrast, Pgk1 and the Hxk1/2 paralogs, which were not significantly enriched in the interactome, were not detected in TRM eluate immunoblots (data not shown). As an additional check on the validity of the interactome data, we determined if Tsa1 interaction with selected targets was dependent on its tagging with MBP. MBP-tagged versions of Pyk1, Nce103, and Fba1 were constructed and functionally verified by complementation (data not shown). The interaction of myc-tagged Tsa1 with each fusion was examined by MBP purification and immunoblotting (**Supplemental Fig. S8A**). Wild-type Tsa1 interacted with all three targets, and the C171S mutation enhanced the interactions. These observations validated the reciprocal results and confirmed that Tsa1-target interactions were not an artifact of Tsa1 modification with MBP.

**Figure 7.**
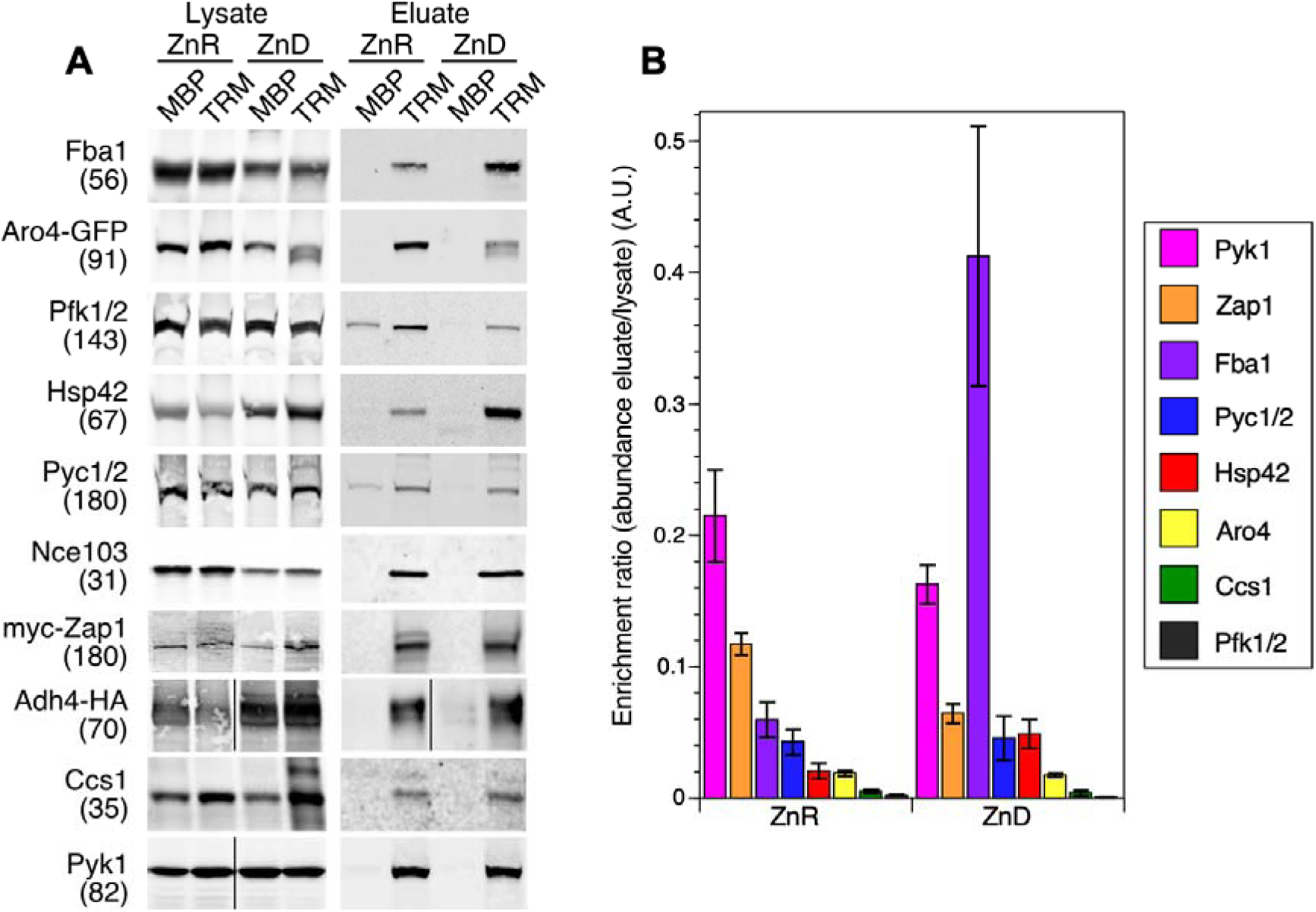
Independent validation of interactome components. A) Wild-type yeast (BY4742) expressing MBP alone, or a congenic *tsa1Δ* mutant expressing Tsa1^C171S^-MBP were grown to log phase in ZnR and ZnD media, cells treated with NEM, and protein extracts prepared with NEM. MBP purification was performed by loading 2 mg input protein on amylose resin columns. Equal volumes of non-reduced column eluates were fractionated by SDS-PAGE, and putative interacting proteins were detected by immunoblotting with specific antibodies. For myc-Zap1, Aro4-GFP, Adh4-HA and FLAG-Nce103, experiments were performed using wild-type (BY4742) or congenic *tsa1Δ* derivatives expressing tagged proteins and detected with anti-tag antibodies. Aro4-GFP was expressed from the genomically tagged locus in BY4742/ARO4-GFP and the *tsa1Δ* derivative CWM355, and Adh4-HA from the tagged locus in ABY64 and the *tsa1Δ* derivative CWM384. FLAG-Nce103 was expressed from the pFLAG-Nce103 plasmid. Transcriptional autoregulation of Zap1 was eliminated by expressing the protein at a low level from a plasmid containing a *GAL1* promoter-driven myc-tagged *ZAP1* gene (pYEF2-myc-Zap1) in glucose-grown cells. Apparent mass of the detected proteins in kDa is indicated in parentheses (left), as determined by comparison with molecular mass markers. B) Relative enrichment of selected interactome components in TRM eluate vs. lysate fractions as determined by immunoblotting. Data was generated by quantitation of three experimental replicates, including the immunoblots shown in A. Values shown are the average ± 1 S.D.

Because MS is sensitive, and the interactome analysis does not factor in relative protein abundance in lysates, even minor components of the TRM eluate might be significantly enriched vs MBP alone. To address this issue, we determined the relative degree of Tsa1 association with selected interactome targets by quantifying immunoblots of input TRM lysate fractions vs eluate fractions (**Fig. 7B**). This analysis revealed that Pyk1 and Zap1 were the most highly enriched in ZnR TRM eluates vs lysates, suggesting they were more tightly associated with Tsa1. This analysis thus provides a means to rank Tsa1 interactions in order of their potential regulatory importance. This experiment also revealed zinc-dependent changes in Tsa1 association, possibly identifying which targets contribute to adaptation. For Fba1, we observed a near 7-fold increase in enrichment in TRM fractions in ZnD samples (**Fig. 7B**). We previously observed that Tsa1 was required for full activation of apo-Fba1 by zinc *in vitro* [20]. The observation that Fba1 and Tsa1 association was enhanced in ZnD cells is consistent with Tsa1 supporting Fba1 activity under these conditions.

### Identification of novel Tsa1 mixed disulfide species

Our observation that both wild-type Tsa1 and Tsa1^C171S^ formed a mixed disulfide with Pyk1 (**Supplemental Fig. S4)** was consistent with a role in the redox regulation of this enzyme (**Fig. 1A**, step 3). Likewise, the identification of mixed disulfides of Tsa1^C171S^ and targets would be consistent with a role for Tsa1 in their regulation. To identify such intermediates, we purified TRM from cells without NEM pretreatment, reasoning that allowing limited oxidation of TRM *in vitro* would enhance formation of mixed disulfides. These samples were examined for the presence of high molecular weight reduction-sensitive species of putative Tsa1 partners co-purifying with TRM. We identified three proteins (Nce103, Fba1, and Zap1) that formed reduction sensitive species with lower gel mobility **(Fig. 8** and **Supplemental Fig. S9**). The abundance of Nce103 and Fba1 in eluates was too low to definitively determine if their redox-sensitive forms contained TRM (multiple TRM bands were present at the expected positions of these species), and it is possible that these species are multimers of the target protein alone, or mixed disulfides formed with other proteins. In the case of Zap1 however, mixed disulfide species were clearly identifiable. In purified TRM fractions, myc-tagged Zap1 was substantially enriched when compared to MBP alone fractions (compare **Fig. 8A, B**). Reduced myc-tagged Zap1 migrated just below the 180 kDa molecular marker (**Fig. 8B**, *right* panel). In non-reduced samples, an additional reduction-sensitive band with higher apparent molecular weight was detected (**Fig. 8B**, *left* panel). In contrast, when NEM pretreatment was omitted, Zap1 was detected as several species migrating above 180 kDa (**Fig. 8C**, *red* channel in overlay). These also contained TRM (**Fig. 8C**, *green* channel in overlay), indicating they represented mixed disulfides and potential intermediates in a redox relay process. In comparison to other Tsa1 targets such as Pyk1, the abundance of the oxidized forms of Zap1 relative to the reduced form was very high (compare **Fig. 8C** with **Supplemental Figure S9**), even when cells were NEM treated prior to fractionation (**Fig. 8B)**. This observation suggested that Zap1 was particularly sensitive to oxidation by Tsa1.

**Figure 8.**
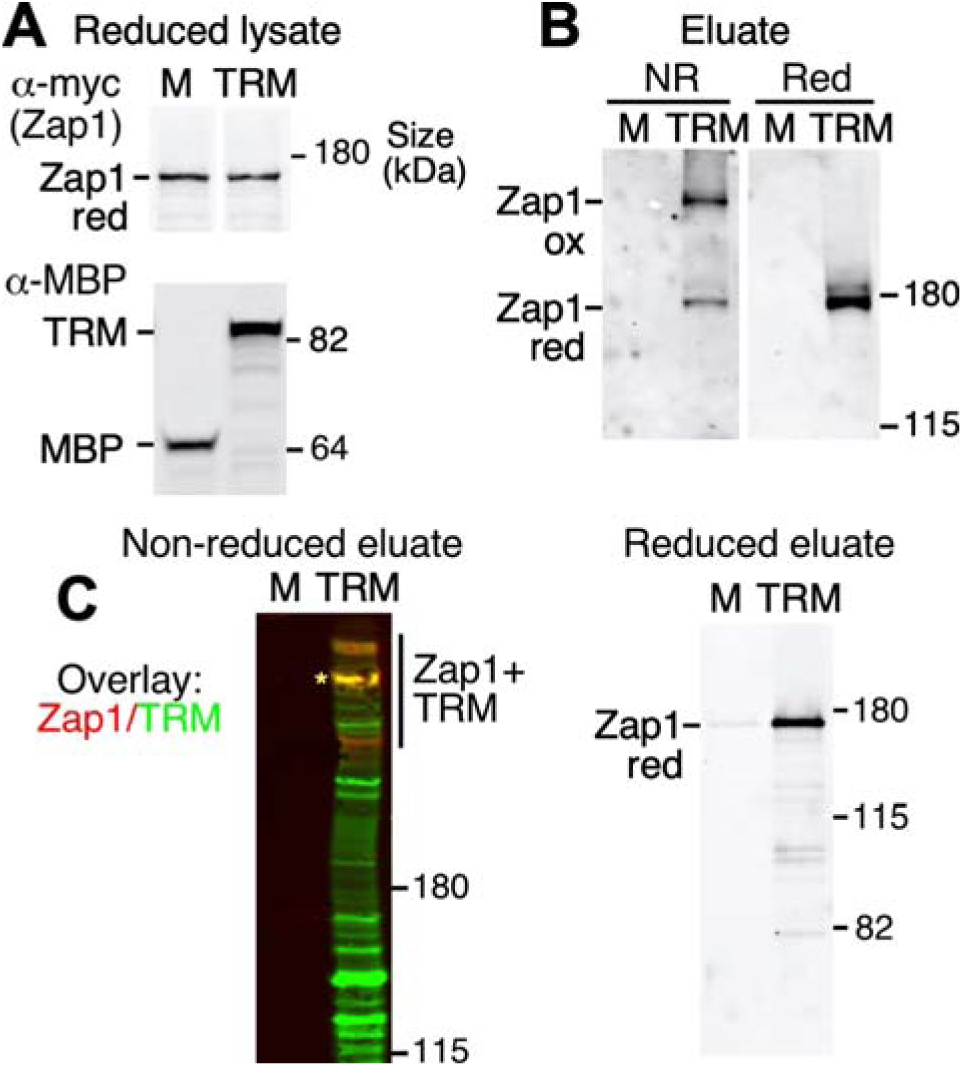
Tsa1^C171S^ interacts with and oxidatively crosslinks to full-length Zap1. A) A yeast strain expressing C-terminally myc-tagged Zap1 (Zap1-13xmyc) was transformed with pMBP (M), and a congenic *tsa1Δ* mutant (CWM347) was transformed with pTsa1^C171S^-MBP (TRM). Strains were grown to log phase in ZnR medium and cells treated with 5 mM NEM before protein extraction. Reduced protein samples were subject to SDS-PAGE and immunoblotting to detect MBP and myc-tagged Zap1. B) MBP was purified from cell lysates shown in (A) and reduced with DTT (*right*) or not treated (*left*). Zap1 was detected by immunoblotting with anti-myc. C) TRM or MBP was purified from ZnR cells as described in A, except that cells were not pretreated with NEM. MBP or TRM (*green*) and Zap1 (*red*) were detected by immunoblotting and their signals overlain. The *asterisk* indicates a major species that reacted strongly with both antibodies. In contrast to the results shown in panel B, the anti-myc antibody did not detect a band at the size expected for monomeric Zap1 (∼180 kDa), but this band was restored upon sample reduction (C, *right* panel). Minor bands below reduced Zap1 are degradation products.

The identification of Zap1 as a target for Tsa1 oxidation was exciting because it implied that the severe growth defect of ZnD *tsa1Δ* cells might be due to the loss of a Tsa1-dependent function of Zap1. Zap1 contains discrete functional domains with well-established roles in zinc-responsive transcriptional regulation, any of which might be a target for Tsa1. The C-terminal DNA binding domain (DBD) contains five zinc finger motifs (ZnF3-7) (**Fig. 9A**, black bars) that allow Zap1 to bind Zn responsive elements (ZRE) in the promoters of its target genes [73, 74]. The activity of the DBD is independently regulated by Zn status [75] by an as yet unknown mechanism. Two other centrally located Zn finger motifs (ZnF1 and 2) form part of the AD2 activation domain [76]. Zinc binding by these ’labile’ zinc fingers inhibits AD2 activity. The activity of a second N-terminal activation domain (AD1) is also Zn responsive and likely directly regulated by Zn binding [77].

**Figure 9.**
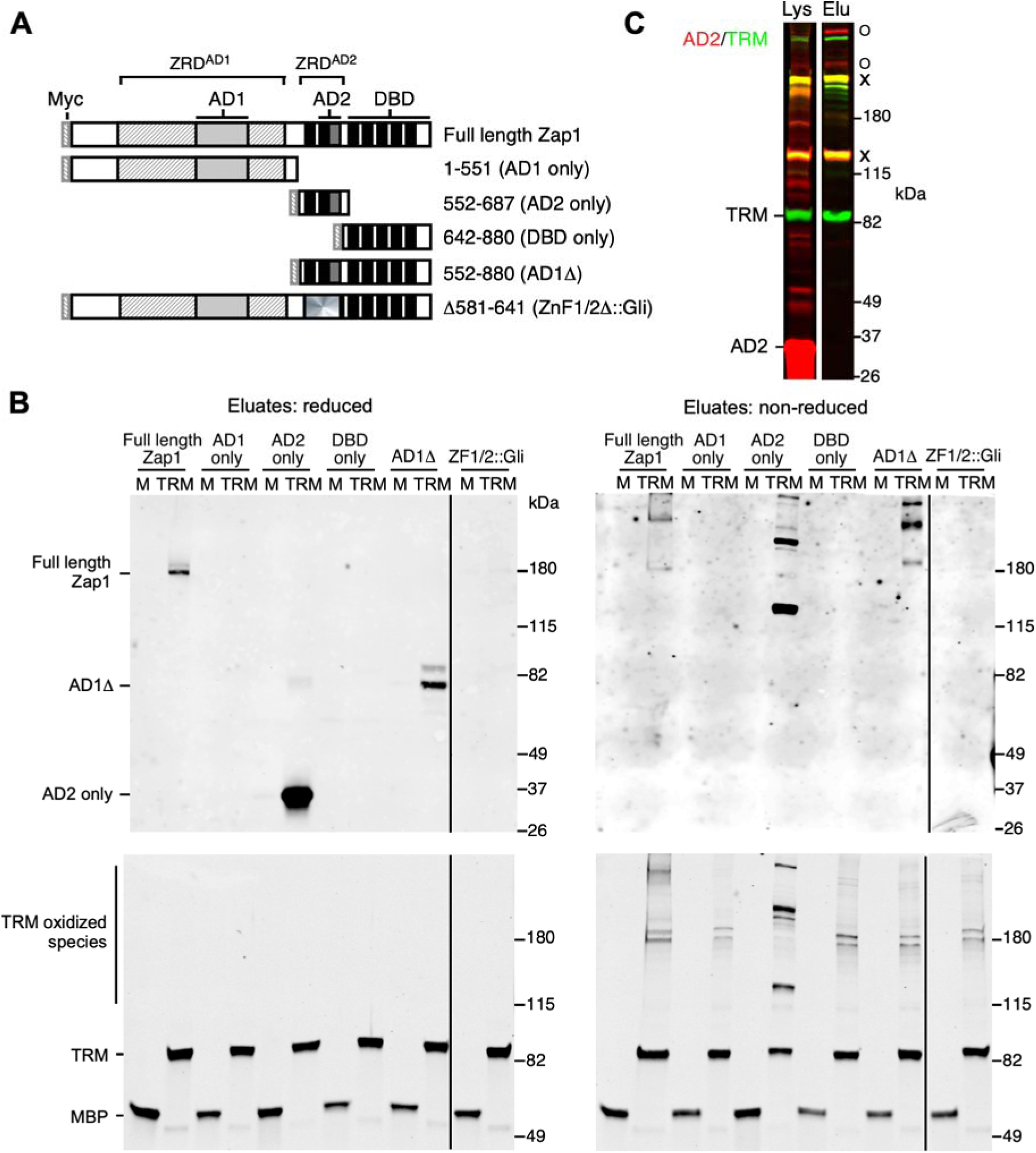
Tsa1^C171S^ interacts with and oxidatively crosslinks to Zap1 AD2. A) Domain structure of N-terminally myc-tagged Zap1 plasmid and its derivatives used in these experiments. The plasmids used were pYEF2-myc-Zap1 (full length Zap1), pYEF2-myc-Zap1^1-551^ (AD1 only), pAFH13 (AD2 only), pAB13 (DBD only), pYEF2-ZAP1^552-880^ (AD1Δ), and pYEF2-myc-Zap1^Znf1/2::GliZnf1/2^ (ZnF1/2Δ::Gli) (see **Supplemental Table S7**). In the ZnF1/2Δ::Gli plasmid, the Zap1 AD2 activation domain (residues 582-640) is replaced with an homologous 58 amino acid zinc finger pair from the Gli transcription factor (*radial shading*). This construct thus retains the general structure of full length Zap1 but lacks one of the zinc-responsive activation domains. B) Wild-type BY4742 and a congenic *tsa1Δ* mutant were transformed with pMBP (M) or pTsa1^C171S^-MBP (TRM) and additional plasmids to express the indicated Zap1 derivatives. Zap1 and most derivatives were expressed from the *GAL1* promoter by growth in ZnR medium containing 2% glucose and 1% galactose, which generated a moderate level of expression. The AD2 only construct was expressed from the constitutive *PGK1* promoter. Cells were treated with NEM and protein extracted for MBP purification. Samples were reduced (*left* panel) or left untreated (*right* panel) before SDS-PAGE and immunoblotting. Zap1 was detected with anti-myc, and MBP or TRM with anti-MBP. C) Overlay comparison of non-reduced samples of lysates (*left*) and eluates (*right*) expressing Zap1 AD2 only (*red* channel) and TRM (*green* channel). *Yellow* bands marked with “X” are oxidized species reacting with both antibodies, and *red* bands marked with “O” are oxidized Zap1 species lacking a TRM component.

To identify the Tsa1 interaction site, we used a set of myc-tagged derivatives of Zap1 missing one or more major domains (**Fig. 9A**), and co-expressed these with TRM to localize the region(s) required for Zap1 oxidation. Since there was little effect of Zn deficiency on the Tsa1-Zap1 interaction (**Fig. 7B**), Zap1 derivatives were expressed from the *GAL1* or *PGK1* promoter in ZnR cells. Derivatives accumulated to similar levels **(Supplemental Fig. S10)**, with the exception of “AD1 only”, which was lower abundance but still detectable. After purification, only those derivatives containing the AD2 domain co-fractionated with TRM (full length myc-Zap1, AD2 only, and AD1Δ) (**Fig. 9B)**. Analysis of non-reduced eluate samples revealed multiple slowly migrating reduction-sensitive species of all three of these derivatives, indicating substantial oxidation. In the case of “AD2 only”, some oxidized species also contained TRM, as indicated by their reaction with both antibodies (**Fig. 9C**, *yellow* bands, *right* lane). Surprisingly, these species were also highly abundant in unfractionated lysate samples (**Fig. 9C**, *yellow* bands marked with “X”, *left* lane), indicating AD2 was efficiently oxidized by Tsa1^C171S^. In addition to these species, we also detected oxidized AD2 species that co-purified with TRM but were not crosslinked to it, as indicated by their absence from the MBP (green) channel in overlain immunoblots (**Fig. 9C**, *red* bands marked with “O” in *right* lane). These species were also detected in lysates (**Fig. 9C**, *left* lane). Their lower mobility in SDS-PAGE is consistent with the formation of crosslinked AD2 homo-multimers, or mixed disulfide species formed with unknown proteins. This is an important observation because these species likely represent products of disulfide exchange by TRM-AD2 mixed disulfides, as predicted by the redox relay model (**Fig. 1A**, step 4). Notably, the ZnF1/2Δ-Gli construct, in which Zap1 ZnF1 and 2 are replaced with an homologous domain of the Gli protein, delineated the region required for AD2 oxidation to Zap1 residues 582-640 (**Fig. 9B**). This region contains only six cysteine residues (Cys 585, 590, 618, 623, 631 and 639). Interestingly, three of these (586, 618, and 623) are Zn ligands located in the two labile Zn fingers of AD2. Their oxidation would be expected to inhibit zinc binding by these motifs, possibly resulting in AD2 activation in the presence of free zinc. In summary, we found that the AD2 domain of Zap1 was both necessary and sufficient for its oxidation by TRM ,and that oxidation of this domain is followed by disulfide exchange and the release of oxidized AD2. In addition, our analysis suggests that by preventing Zn from binding the AD2 zinc finger motifs, Tsa1 could increase AD2 activity in Zn-replete cells.

### Peroxide-induced formation of Zap1-Tsa1 mixed disulfide species

The detection of Zap1 AD2 and TRM mixed disulfide species in purified fractions implied that Zap1 was oxidized *in vivo*. However, as it was possible that the oxidation we observed occurred *in vitro* during sample fractionation, we also examined Zap1 oxidation in samples prepared using a method that completely preserves redox state. Cells expressing both untagged Tsa1^C171S^ and the AD2 domain of Zap1 tagged with a C-terminal fragment of GFP (VC) were treated (or not treated) with hydrogen peroxide, then treated with trichloroacetic acid (TCA) to fix the redox state of cysteines prior to protein extraction. Reduced or unreduced protein samples were then subject to immunoblotting with anti-Tsa1 and anti-GFP (VC) antibodies. These experiments (**Fig. 10A**) confirmed that in response to H_2_O_2_ treatment, cells expressing both Tsa1^C171S^ and VC-AD2 accumulated a mixed disulfide species of approximately 70 kDa containing both proteins. This observation indicated both that this reaction occurred *in vivo,* and that it was not an artifact of Tsa1 modification.

**Figure 10.**
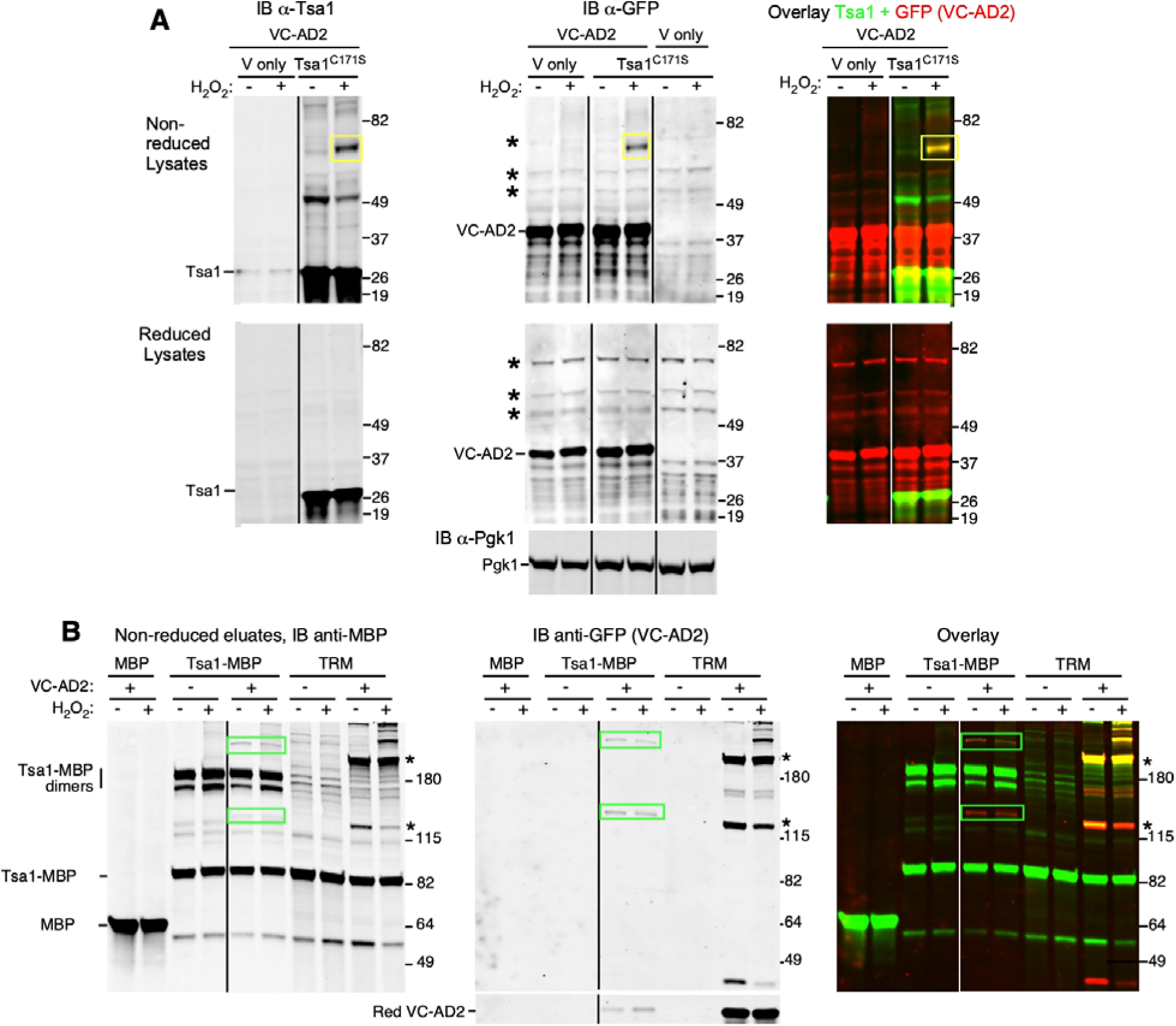
Tsa1 oxidizes Zap1 AD2 *in vivo*. A) Cultures of a *tsa1Δ* strain transformed with pVC-AD2 (Zap1 AD2 fused to the C-terminal half of GFP) and either empty vector (pFL38) or pTsa1^C171S^ were grown in zinc-replete medium to log phase. Cells were treated with 0.5 mM peroxide for 10 min, then with TCA to fix redox state (final concentration 12%). Protein was extracted with TCA, and samples were reduced with DTT (*lower* panels) or not treated (*upper* panels) prior to SDS-PAGE and immunoblotting with anti-Tsa1 (*left* panels) or anti-GFP (for VC-Zap1AD2, *center* panels). Pgk1 was also detected as a loading control (IB α-Pgk1). Note minor band detected at the Tsa1 position in *upper left* panel is Tsa2. Color panels (*right*) show overlay of AD2 and Tsa1 images. *Yellow* boxes indicate the position of a reduction-sensitive species composed of both proteins. Asterisks indicate major species reacting non-specifically with anti-GFP. B) Wild-type Tsa1-MBP forms mixed disulfides with Zap1 AD2. A wild-type strain (BY4742) expressing MBP and VC-AD2, or a congenic *tsa1Δ* mutant strain expressing wild-type Tsa1-MBP or TRM ± VC-AD2 from plasmids as indicated, were grown to log phase in ZnR medium and treated with 10 mM NEM. Prior to NEM treatment, some cultures were treated with H_2_O_2_ as indicated (0.5 mM for 10 min). Cell lysates were prepared, MBP purified, and non-reduced samples subject to SDS-PAGE and immunoblotting to detect MBP (*left* panel) or GFP (VC-AD2, *center* panel). Major reduction-sensitive species containing both Tsa1-MBP (or TRM) and VC-AD2 are indicated in overlain blots (*yellow* bands marked with *asterisks* for TRM, *green* boxes for wild-type Tsa1-MBP). For all immunoblots, *black* (individual blots) or *white* lines (overlays) indicate where superfluous lanes were removed. For B, reduced samples were also analyzed in parallel (see **Supplemental Figure S11**).

It was also formally possible that the formation of the TRM-AD2 mixed disulfide was an artifact of the C171S mutation. To determine if wild-type Tsa1 could also oxidize AD2, we examined native cell lysates and purified MBP fractions after H_2_O_2_ treatment of cells expressing wild-type Tsa1-MBP, TRM and VC-AD2 (**Fig. 10B**). In MBP fractions purified from cells expressing TRM, we detected VC-AD2 below the 49 kDa marker (*center* panel, anti-GFP), as well as two very abundant forms with lower mobility (>115 and 180 kDa, *asterisks*). These species also reacted with MBP antibody (**Fig. 10B**, *left* panel and overlay) and were eliminated by reduction (**Supplemental Fig. S11**), indicating they were mixed disulfides formed with TRM. Reduction sensitive species with similar mobility were also seen in lysates of cells expressing wild-type Tsa1-MBP (**Fig. 10B**, anti-MBP, *green* boxes). These species were also mixed disulfides as they were detected by both antibodies (**Fig. 10B**, overlay, *green* boxes). In purified TRM fractions, H_2_O_2_ treatment resulted in the depletion of VC-AD2 and the appearance of new redox-sensitive species of >180 kDa, which also contained TRM (**Fig. 10B**, overlay). In comparison, only two mixed disulfide species of VC-AD2 were detected in wild-type Tsa1-MBP fractions, equivalent in size to the two major forms detected in TRM fractions. VC-AD2 was completely crosslinked to Tsa1-MBP even in the absence of H_2_O_2_ treatment, meaning it was unclear if these intermediates formed *in vivo*. However, this experiment confirmed that wild-type Tsa1 can target the AD2 domain for oxidation and form the same mixed disulfide species as Tsa1^C171S^.

### Tsa1 is required for maximal Zap1 activity in severely zinc-deficient cells

If the purpose of the Tsa1-mediated oxidation of Zap1 is to regulate Zap1 activity, we expected that the loss of Tsa1 would affect Zap1 function. To test this prediction, we examined the effect of the *tsa1Δ* deletion mutation on the expression of five well-characterized Zap1 target genes. Wild-type and *tsa1Δ* cells were grown in ZnR and ZnD medium for 48 hours and RNA was extracted for RT-qPCR to measure target mRNA levels. Relative gene expression was calculated by normalizing each to a set of three control mRNA known to be unaffected by zinc status (*ACT1*, *TAF10* and *CMD1*) [8]. The results (**Fig. 11A**) showed that the *tsa1Δ* mutation had little effect on gene expression in ZnR cells, but substantially reduced target expression in ZnD cells. Because *tsa1Δ* cells grow more slowly than wild-type in low zinc (**Fig. 11B**), part of this effect might be explained by the lower zinc requirement of slow- vs. fast-growing cells, which could downregulate Zap1 activity. To investigate this possibility, we took advantage of our recent discovery that supplementing ZnD cells with PEP greatly increased their growth rate [20]. Supplementing *tsa1Δ* ZnD cells with PEP restored their growth rate to that of wild-type ZnD cells, thus eliminating growth rate as a confounding factor in Zap1 activity. While this intervention increased the expression of *ZRT1*, *ZAP1*, and *ZRT3* in ZnD *tsa1Δ* mutant cells, it did not restore wild-type levels of expression to any of the six genes (**Fig. 11A**). Thus, the failure of ZnD *tsa1Δ* cells to fully induce Zap1 target gene expression is not due solely to their slow growth, and is consistent with intrinsically lower Zap1 activity.

**Figure 11.**
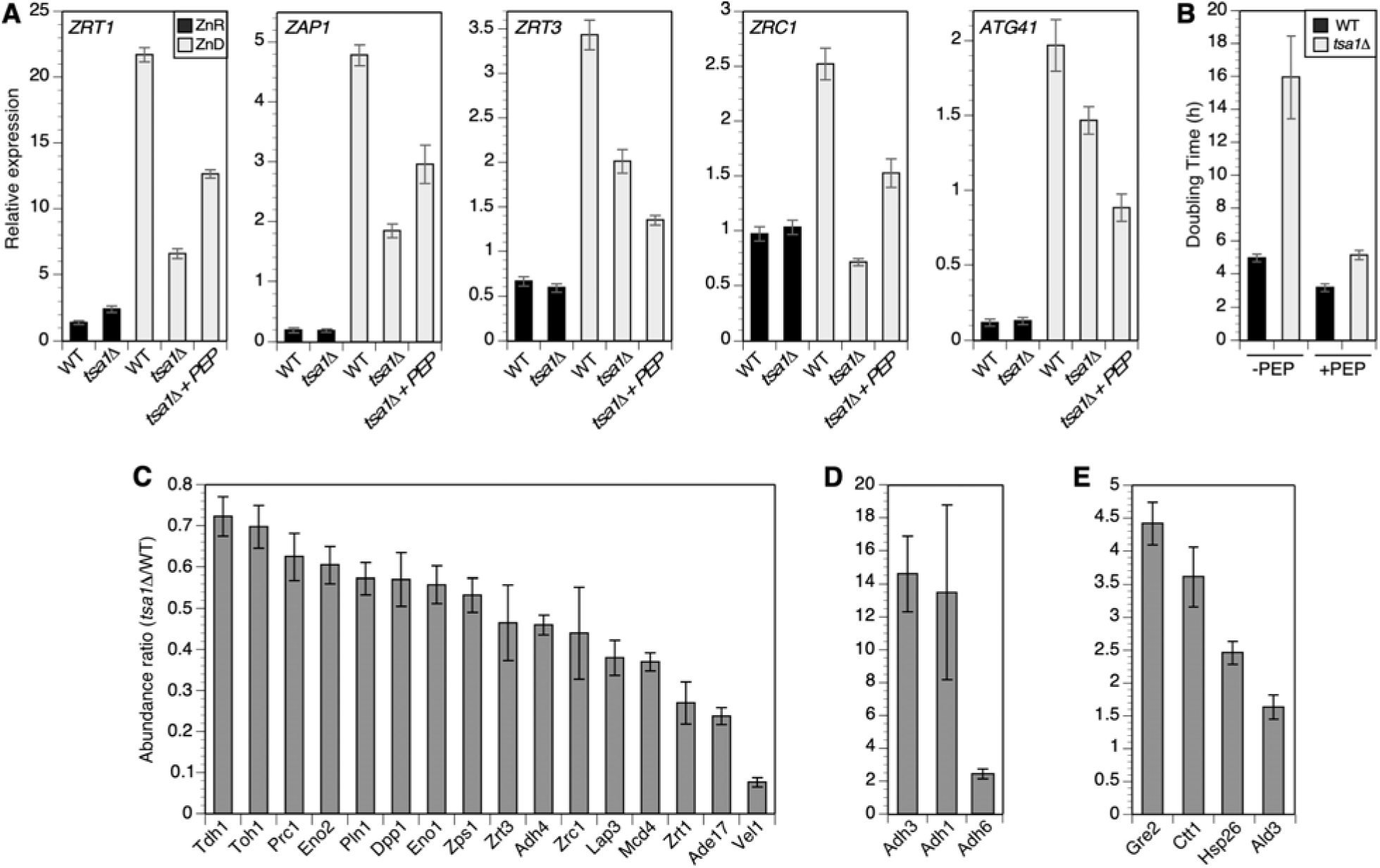
Tsa1 supports Zap1 activity in ZnD cells. A) Wild-type (BY4742) or congenic *tsa1Δ* mutant cultures were grown in SC medium and used to inoculate aliquots of LZM + 0.5 μM ZnCl_2_ to an initial A_595_ of 0.05 (for wild type) and 0.1 (for *tsa1Δ*). Cultures were maintained at < 0.3 *A*_595_ by dilution into fresh medium, then harvested after 48-hours and RNA extracted for qRT-PCR analysis. The relative abundance of each of the five Zap1 target gene mRNAs was normalized to three control mRNA (*ACT1*, *TAF10* and *CMD1*), as described in Experimental Procedures. Values are the average of five replicates and error bars indicate ± 1 S.D. B) Effect of 30 mM PEP supplementation on the doubling time of wild-type and *tsa1Δ* mutant cells in ZnD medium (LZM + 0.5 μM ZnCl_2_). ZnD cultures were inoculated and maintained as described for panel A. Doubling time was measured over a 30-hour period from 14 - 44 h of culture. Values are the average of five replicates and error bars indicate ± 1 S.D. C-E) Relative abundance of Zap1-regulated proteins in a *tsa1Δ* mutant (CWM254) vs wild-type (BY4742/*arg4Δ*) cells after growth in low zinc (LZM + 0.7 μM Zn). Ratiometric quantitation of the indicated proteins in WT and *tsa1Δ* strains was performed using SILAC (see Experimental Procedures). Abundance ratio values represent *tsa1Δ* samples normalized to wild type (four experimental replicates, average ± 1 S.D.).

We then performed a proteomic survey to measure the accumulation of Zap1-regulated proteins in wild-type vs *tsa1Δ* strains grown for an extended period (48 h) in low zinc. Quantitative comparison of protein abundance was achieved via metabolic labeling and GC-MS/MS (see Experimental Procedures). Consistent with the RT-qPCR data, we observed a substantial decrease in the accumulation of the majority of Zap1-regulated proteins **(Fig. 11C**). The exceptions included those for which expression is normally repressed by Zap1 in ZnD conditions (*e.g., ADH1* and *ADH3*). Accumulation of these proteins was elevated in ZnD *tsa1Δ* cells, consistent with less effective gene repression by Zap1 **(Fig. 11D**). Four more Zap1-activated targets showed elevated abundance in *tsa1Δ* relative to wild-type **(Fig. 11E**). However, the genes encoding those proteins are known targets of stress-responsive regulatory factors such as Msn2/4 and Hsf1, and their induction likely reflects stress experienced by the ZnD *tsa1Δ* mutant [9]. In summary, these experiments indicated that Tsa1 is required for the full activity of Zap1 under conditions where zinc supply is restricted for a sustained period of time. Given the importance of Zap1 for survival in low zinc, the contribution of Tsa1 to Zap1 function may contribute to the severe growth defect seen in ZnD *tsa1Δ* cells.

## DISCUSSION

In this report, we describe a new approach to characterizing potential regulatory interactions of the 2-Cys PR Tsa1, which resulted in the identification of numerous novel interacting proteins. This enquiry was prompted by our discovery that the redox sensitive glycolytic enzyme Pyk1 was downregulated by Tsa1 in ZnD cells to conserve the glycolytic intermediate PEP and maintain aromatic amino acid synthesis [20]. Consequently, we suspected that Tsa1’s redox sensor function was involved in adapting glycolysis and possibly other processes to low zinc. Our major goals for this work were to determine the importance of the redox sensor function of Tsa1 for adaptation to low zinc, and to identify novel Tsa1-regulated proteins that might contribute to this adaptation. This report provides both important general insights into PR function, and specific insights into the role of Tsa1 in the adaptation to zinc deficiency.

### Role of Tsa1 in adaptation to ZnD conditions

Our working hypothesis at the beginning of this study was that Tsa1 chaperone activity was essential for growth in low zinc, and that this activity stabilized zinc apoproteins and thus prevented the deleterious accumulation of misfolded proteins [9]. Subsequent work has supported some of the major predictions of this model. For example, we showed that zinc-deficient cells accumulated large amounts of potentially unstable zinc apoproteins, and we also observed a disproportionate depletion of zinc-dependent proteins upon induction of zinc deficiency, consistent with their enhanced degradation [15]. In addition, we discovered a role for autophagy in the adaptation to low zinc [78, 79] and found that the *UBI4* ubiquitin gene was both essential for growth in low zinc [79], and upregulated by Zap1 to maintain fitness [8]. Together this work showed that mechanisms of protein homeostasis are important for adaptation to low zinc. However, a direct role for Tsa1 in protein homeostasis in ZnD cells proved more difficult to support. Because zinc plays a role in stabilizing protein structure, we proposed that excessive misfolding and aggregation of zinc apoproteins contributed to the *tsa1Δ* growth defect. However, we were unable to confirm this prediction. For example, a mass spectrometric survey of purified protein aggregates from zinc-deficient *tsa1Δ* mutant cells showed that contrary to expectations, zinc-dependent proteins did not disproportionately accumulate in detergent-resistant protein aggregates (data not shown). We also surveyed GFP-tagged alleles of the 100 most abundant zinc-dependent proteins to identify any that specifically formed visible aggregates in ZnD *tsa1Δ* cells [9]. Although we did identify several such proteins, further study revealed that their aggregation was dependent on GFP addition (data not shown), indicating their instability was likely artifactual. Thus although we cannot exclude the possibility that the Tsa1 chaperone function makes some contribution to protein stability in low zinc, we have not yet found direct evidence to support this hypothesis.

In contrast, this report and our previous work [20] argue for the importance of Tsa1 redox sensor function to fitness of ZnD yeast. First, our examination of additional Tsa1 mutants indicated that the redox sensor function of Tsa1, and not its chaperone activity, is essential for growth in low zinc (**Fig. 1B**). Second, we obtained additional evidence that Pyk1 is a target for redox regulation by Tsa1. In a previous study, we showed that Pyk1 activity was redox sensitive, and that Tsa1 inhibited Pyk1 activity in ZnD cells. These and other observations suggested that Tsa1 facilitated adaptation to zinc deficiency by inhibiting Pyk1 activity in order to conserve its substrate PEP for aromatic amino acid synthesis. Consistent with a redox relay model for Pyk1 regulation, we now show that Tsa1 and Pyk1 interact non-covalently *in vivo*, and that their interaction was redox dependent. Further support for a redox relay comes from the observation that wild-type Tsa1 formed a mixed disulfide with Pyk1 in response to H_2_O_2_ *in vivo*, and that the formation of this intermediate was dependent on a specific Pyk1 residue (C174). Although the mechanism of Pyk1 redox regulation and the specific role of C174 require further investigation, these observations provide further support for Tsa1 acting as a redox sensor for Pyk1, which may explain the importance of Tsa1 for Pyk1 downregulation in ZnD cells [20]. One question that remains unanswered is if formation of the mixed disulfide of Pyk1 and Tsa1 is followed by disulfide exchange to produce oxidized Pyk1 and reduced Tsa1 (**Fig. 1B**, step 4). Although it is possible that this process occurs, it is also possible that formation of the Pyk1-Tsa1 mixed disulfide is the final regulatory step in this process, as the location of C174 in the Pyk1 B-domain suggests crosslinking to Tsa1 would be sufficient to inhibit its activity. To restore basal Pyk1 activity (**Fig. 1B**, step 5), this crosslink could be reversed by disulfide exchange with Tsa1 C171, or reduced by thioredoxin.

### Functional verification of Tsa1-MBP

A major goal of this work was to identify novel Tsa1-interacting proteins, particularly those that might impact tolerance to low zinc. The redox sensor model predicts that Tsa1 physically associates with its regulatory targets prior to the generation of a redox signal, suggesting these proteins would copurify with Tsa1. However, previous studies identified relatively few proteins interacting with Tsa1 or other PR’s. Working from the hypothesis that Tsa1-target interactions would be redox sensitive (*i.e.,* inhibited by oxidation), we developed a system to allow rapid and direct purification of Tsa1 and associated proteins from cell lysates while suppressing their oxidation with NEM. A Tsa1-MBP fusion was constructed and extensively tested to verify that it identified genuine Tsa1 interactions. For example, we verified that tagging Tsa1 with MBP did not affect its stability, an important consideration given previous work [36, 66], and our observations (**Supplemental Fig. S2B**), that Tsa1 C-terminally tagged with GFP aggregated *in vivo*. This behavior was an artifact of tagging, as the GFP tag reduced Tsa1 solubility in ZnD cells (**Supplemental Fig. S2C, D**), and co-expression with untagged Tsa1 eliminated Tsa1-GFP aggregation into intracellular foci (**Supplemental Fig. S2B**). Given that Tsa1 assembles into decamers [80], the simplest interpretation of this observation is that the association of stable untagged Tsa1 with an unstable tagged form suppressed its instability. The possible effect of Tsa1 C-terminal additions on stability is a major concern for studies of interaction, as aggregation might promote artifactual Tsa1 association with protein chaperones or other misfolded proteins. Several lines of evidence argue that Tsa1-MBP retained the full function of untagged Tsa1: first, Tsa1-MBP solubility was little different from untagged Tsa1; second, Tsa1-MBP effectively complemented the *tsa1Δ* mutant growth defect in low zinc; and third, the interactome included only three protein chaperones (notably, it did not include Hsp104, which associates with aggregates of unfolded proteins) [36]. Lastly, we independently confirmed that myc-tagged Tsa1 also associated non-covalently with MBP-tagged interactome components (**Fig. 7**), indicating that these interactions were not artifacts of tagging Tsa1 with MBP. Thus, we extensively verified the suitability of Tsa1-MBP as a tool to identify novel interacting proteins. We suspect that PR-MBP fusions may also prove useful for identifying and studying PR interactions in higher eukaryotes.

### Tsa1-target interactions are predominantly non-covalent

Using the Tsa1-MBP fusion, we proceeded to characterize Tsa1’s interaction with Pyk1, a known regulatory target. An interaction of Pyk1 and Tsa1 was previously detected by immunoprecipitation, and was dependent on Tsa1 C48 [61]. This dependence could reflect covalent association by disulfide crosslinking, or non-covalent interaction via the Tsa1 chaperone activity (which is dependent on C48 hyperoxidation). We made several key observations that differed from this previous report. First, we identified a stable non-covalent interaction that was independent of Tsa1C48. Second, this stable interaction was sensitive to Tsa1 and/or Pyk1 oxidation, as it was enhanced by treatment of cells with the cysteine alkylator NEM prior to cell fractionation, and also by Tsa1 mutations that eliminate or strongly inhibit peroxidase activity (*e.g.,* C48S and C171S), and thus directly prevent or inhibit Tsa1 oxidation. Third, the non-covalent interaction was independent of Tsa1 chaperone activity, as a mutation that blocks that function (C48S) actually enhanced Tsa1’s association with Pyk1. Interestingly, we also observed that NEM treatment of cells enhanced the non-covalent interaction of both Tsa1 cysteine mutants and Pyk1. This observation suggests that in addition to inhibiting Tsa1 oxidation, the effect of NEM is partly due to blocking target oxidation, *i.e.,* the oxidation state of Tsa1 targets may also influence their interaction with Tsa1. This redox sensitive non-covalent binding of targets by Tsa1 has important implications for Tsa1 function in regulation. Redox sensitivity may be a general property of Tsa1 interactions, as treatment with cysteine alkylators enhanced the co-fractionation of multiple proteins with TRM (data not shown). It has been shown for both native and non-native PR targets that proximity of PR’s to targets confers greater efficiency and specificity to redox regulation [40, 50].

### Expansion of the Tsa1 interactome

Previous studies identified a limited number of proteins that copurified with or interacted *in vitro* with Tsa1. The BioGRID database included 88 proteins reported to associate with Tsa1 (**Supplemental Table S3**). These interactions were predominantly detected with bait proteins other than Tsa1, using methods such as affinity capture combined with MS or immunoblot analysis, co-fractionation, proximity labeling, or two-hybrid analysis. In this study, we used a novel purification tag combined with IAA treatment and label-free MS to identify Tsa1^C171S^-associated proteins. The rationale for using the C171S mutant was that by promoting the accumulation of Tsa1-target mixed disulfide species, and by conserving non-covalent interactions, we could enhance the capture of novel Tsa1 partners. Accordingly, our MS analysis identified over 170 Tsa1 partners, of which only 15 were previously reported in BioGRID (**Supplemental Table S3**). To explain this minor overlap with the BioGRID list, we suggest that many large-scale methods may not accurately report Tsa1 interactions for the reasons outlined above, *i.e.* that modification of Tsa1 or other proteins with functional tags can alter their function and stability, or that genuine interactions may be suppressed when Tsa1 redox state is not preserved *in vitro*. For these reasons we believe that our method more accurately reflects the Tsa1 interactome *in vivo*. Indirect evidence for the relevance of our approach comes from the observation that key regulatory enzymes of intermediary metabolism were overrepresented in the interactome (**Table 1** and **Fig. 6**). However, further analysis of individual interactome components is required to validate our results.

### Many interactome components are oxidized and redox regulated

It is now widely appreciated that ROS or RNS can serve as signaling molecules to regulate processes such as glycolysis.

However, there are two major questions in the redox regulation field that remain unanswered: first, how do low concentrations of relatively stable ROS such as H_2_O_2_ effectively oxidize cysteine residues *in vivo*; and second, how does a redox signal lead to the oxidation of specific cysteine residues in regulated proteins? An attractive hypothesis is that redox regulation is mediated by thiol peroxidases that react with ROS or RNS, then transfer their oxidation to specific targets. If Tsa1 acts as such a receptor, the interactome should include examples of proteins that are reversibly oxidized *in vivo*. To address this question, we examined data from nine redox proteomic surveys conducted with *S. cerevisiae* or *S. pombe* [58, 81–88] to identify interactome components with reversibly oxidized cysteines. These studies surveyed oxidation of the yeast proteome in normally growing cells, as well as in response to conditions that induce oxidation, such as ROS treatment [82, 84], chronological aging [83], or mutations inactivating antioxidant enzymes [81, 88]. This analysis found 95 interactome proteins (54%) were previously reported to have oxidized cysteines (**Supplemental Table S2,** panel 3, oxidized targets). Thus the majority of the interactome is detectably oxidized and potentially subject to redox regulation (note that due to the technical limitations of shotgun proteomic analysis, the absence of detectable oxidation for an interactome component does not exclude this possibility) [89]. We also surveyed the literature for redox regulated Tsa1 targets, or those with redox sensitive activity. In addition to Pyk1 [20] and Yap1 [49, 90], these include Ssa1 [91–93], Cdc48 [94], Uba1 [95], Yol057w (DPPIII) [96], Sch9 [97], Fba1 [87], and Hem12 [98]. Little is known about how the latter proteins are specifically oxidized *in vivo*. Their association with the Tsa1 redox sensor may provide a mechanistic explanation.

### Tsa1 may facilitate adaptation to low zinc via the regulation of zinc proteins

We show here that the redox sensor function of Tsa1 is essential for growth of zinc-deficient cells, but dispensable for ZnR cells. Why is this activity so critical for adaptation to low zinc? We have identified at least two Tsa1-mediated regulatory effects that contribute to adaptation, the downregulation of Pyk1 to support aromatic amino acid synthesis [20], and the maintenance of Zap1 activity in severely ZnD cells. Given the severe growth defect of ZnD *tsa1* mutants however, Tsa1 likely makes additional regulatory contributions involving its redox sensor function. One novel Tsa1 target, the vacuolar zinc transporter Zrt3, directly influences Zn homeostasis [14], and its identification suggests Tsa1 is involved in regulating intracellular zinc storage. We also found multiple zinc-dependent proteins in the interactome, many of which contain cysteine-zinc ligands. These include such important enzymes as Adh4 (the major isoform of alcohol dehydrogenase in ZnD cells), Sam4 (Homocysteine methyltransferase), Nce103 (carbonic anhydrase), and Rpo31 (a core subunit of RNA polymerase III). Given that zinc apoproteins accumulate in ZnD cells [15], we speculate that Tsa1 is involved in maintaining the function of one or more critical zinc-dependent proteins. For example, Tsa1 might enhance apoprotein stability by oxidatively crosslinking zinc-binding cysteine residues, thus substituting for the structural stability normally provided by zinc binding. Tsa1 might also stabilize zinc apoproteins via oxidation of non-ligand cysteines. In this regard, we recently identified a possible role for Tsa1 in supporting the activity of Fba1 in low zinc [20]. Fba1 is highly dependent on zinc status because its activity is disrupted by loss of its zinc cofactor [15]. Fba1 activity further decreased in ZnD *tsa1Δ* mutant cells, suggesting that Tsa1 stabilized activity. Interestingly, Tsa1 seemed to enhance the reactivation of apo-Fba1 when it was resupplied with zinc *in vitro* [20]. This work provides further evidence for a regulatory relationship, first, by showing that Fba1 interacts non-covalently with Tsa1; second, that this interaction is substantially increased in zinc-deficient cells; and third, that Tsa1^C171S^ and Fba1 form a mixed disulfide (**Supplemental Figure S9**). We suggest that Tsa1 oxidizes apo-Fba1 in order to preserve its ability to bind zinc, perhaps by stabilizing a favorable conformation of the apoprotein [15]. Interestingly, there is evidence for redox regulation of Fba1 in response to reactive nitrogen species [99].

### Tsa1 oxidizes Zap1 and enhances its activity

Zap1 was highly abundant in TRM eluate fractions, suggesting it is a major Tsa1 target. Because of the importance of Zap1 for zinc-deficient cells, we focused on understanding the possible role of Tsa1 in regulating this protein. We detected abundant mixed disulfide species containing full-length Zap1 and TRM, consistent with a redox relay process. Deletion analysis identified a region of Zap1 AD2 as both necessary and sufficient for oxidation by Tsa1. This oxidation occurred *in vivo*, as it was promoted by H_2_O_2_ treatment of cells prior to cell fractionation. We also detected oxidized Zap1 AD2 without crosslinked Tsa1, suggesting that disulfide exchange had occurred and oxidized AD2 was released (**Fig. 1A**, step 4). Finally, we showed that wild-type Tsa1-MBP also promoted the oxidation of Zap1 AD2, as demonstrated by detection of mixed disulfide species. Together, these observations argue that Zap1 is a bona fide target for a redox relay involving Tsa1.

Consistent with redox regulation, we also found that Tsa1 was required for full Zap1 activity under conditions of severe zinc deficiency. Expression of many Zap1 target genes was reduced in a *tsa1Δ* mutant, and this reduction was not a consequence of a slower growth rate affecting zinc requirement. Although we have not yet determined how oxidation of Zap1 AD2 could increase its activity, there are some obvious possibilities. For example, AD2 may be oxidatively crosslinked to other components of the transcriptional machinery, such as the SWI/SNF, SAGA, or Mediator coactivator complexes recruited by Zap1 [100], thus stabilizing the transcription factor assemblies that form at Zap1 target gene promoters. The reduction sensitive AD2-containing complexes lacking Tsa1 that we detected may contain such coactivator proteins.

Alternatively, the oxidation of cysteines in AD2 might activate Zap1 by blocking zinc binding. Inhibition of activation domain function by zinc binding probably occurs to some degree even in severely zinc-deficient cells, and oxidation of cysteine zinc ligands might block this residual activity. Other explanations are possible and require further investigation.

The identification of Zap1 as a Tsa1 target protein also raises the question why, when the zinc sensor domains of Zap1 mediate such an effective transcriptional response to low zinc, yeast evolved a secondary pathway for Zap1 regulation. A possible answer is that redox signals carry information about the environment that could influence cells ability to tolerate a specific combination of Zn deficiency and other environmental challenges. For example, our work suggests that Tsa1 redox regulation of Zap1 is mediated through AD2 alone, and this domain is known to preferentially activate a subset of Zap1 target genes [101], suggesting that a redox signal acting via Tsa1 would promote a more selective Zap1 transcriptional response. This response might more effectively combat a combination of low zinc and elevated ROS, for example when ZnD cells encounter excess heat [102], ethanol toxicity [103], or other stress. We also showed that although AD1 alone is capable of driving the expression of most Zap1 target genes, its activity was inhibited by heat stress, while AD2 was little affected. Given our observations here, an attractive model is that heat-induced ROS triggers Tsa1 to oxidize AD2 Cys Zn ligands and prevent binding. If this modification activates AD2 it might compensate for the inhibition of AD1 activity by heat. Other explanations for redox regulation of Zap1 are possible: for example, if Zn deficiency depletes Zn-dependent components of the transcriptional machinery [104], a stress induced redox signal could activate AD2 to sustain expression of the Zap1 regulon. Lastly, it is possible that Zn deficiency itself triggers a ROS signal [21], contributing to the adaptive response via regulation of Zap1 and other redox-sensitive factors such as Pyk1. Whatever the answer, Tsa1 clearly serves as an important amplifier of Zap1-mediated gene regulation in Zn-deficient cells.

## EXPERIMENTAL PROCEDURES

### Yeast culture and general methods

Yeast strains were routinely grown in rich medium (YP with 2% glucose or 3% glycerol+2% ethanol), synthetic defined (SD) medium, or low Zn medium (LZM) as described previously [15]. LZM lacks added Zn and contains chelators (EDTA and citrate) to reduce Zn availability. LZM was supplemented with ZnCl_2_ (usually 1 or 100 μM) to generate zinc-deficient or replete conditions respectively. To aid growth of S288c-derived strains, LZM was routinely supplemented with a subset of amino acids and inositol as previously described [20]. Fluorescence microscopy to detect GFP, and quantitative reverse-transcriptase-PCR (q-RT-PCR) was performed as previously described [9, 15]. Sequences of the oligonucleotides used for q-RT-PCR are listed in **Supplemental Table S6**.

### Plasmids

Plasmids used in this work are listed in **Supplemental Table S7**. The pRS315-TSA1, pRS315-TSA1^C48D^, pRS315-TSA1^ΔCt^, and pRS313-HA-PYK1 plasmids were provided by Shusuke Kuge (for all plasmids, the indicated location of mutations and deletions are relative to the first methionine of the unmodified protein). The Gal1/10 His6-MBP TEV Ura (12URA-C) plasmid was a gift from Scott Gradia via Addgene (Addgene plasmid # 48305). The pFA6a-13Myc-kanMX6 plasmid was a gift from Jurg Bahler and John Pringle (Addgene plasmid # 39294). All new plasmids were constructed by PCR and homologous recombination in yeast and verified by sequencing. The seven *CDC19* plasmids with individual cysteine to serine mutations were constructed from pCDC19 via mutagenic PCR and recombination in yeast. Mutations were verified by sequencing and their effect on *CDC19* function was determined by complementation of the *cdc19::KanMX4* (CWM307) growth defect on glucose medium (all 7 mutant alleles complemented, data not shown). All plasmids used in this work and their complete annotated sequences are available by request to CWM.

### Yeast strain construction

Yeast strains used in this work are listed in **Supplemental Table S8**. All yeast strains are isogenic to the haploid BY4741 and BY4742 strains [105]. Yeast strains from the diploid BY4743 deletion mutant collection, haploid BY4742 viable deletion mutant collection [106], or GFP-tagged BY4742 collection [107] were obtained from Invitrogen. CWM231 (TEF2pr-GFP) is haploid progeny of a BY4743-TEF2pr-GFP diploid [9]. The ZAP1-13xmyc strain was constructed by amplification of the 13xmyc-*ADH1* terminator-*KanMX6* cassette from the pFA6a-13Myc-kanMX6 plasmid using oligonucleotides that added homology to the Zap1 C-terminus, followed by transformation of BY4741 with the amplicon. The additional *tsa1* mutant strains CWM254, 347, 355, and 384 were generated by amplification of *tsa1::KanMX4* or *LEU2* knockout alleles from existing mutant strains [9] and transformation of target strains (BY4742 *arg4*, Zap1-13xmyc, Aro4-GFP, and ABY64 respectively) selecting for *LEU2* or *KanMX4*-positive clones. The CWM357, 359, 361, 363, 365, 367, 442 and 443 strains with *cdc19* cysteine-serine mutations and a *tsa1* deletion were generated in several steps. Mutant *CDC19* alleles carried in plasmids (**Supplemental Table S7**) were amplified by PCR, and the amplicons used to reintroduce *CDC19* in the *cdc19::KanMX4* deletion mutant CWM307, selecting for restoration of growth on glucose after transformation as previously described [20]. Introduction of the mutations was verified by PCR and sequencing. A *tsa1::HptMX6* allele was then constructed by amplification of *HptMX6* from the pAG32 plasmid [108] to generate an amplicon with flanking homology to *KanMX4/HptMX6* regulatory sequences. This product was used to replace *KanMX4* in BY4742 *tsa1::KanMX4* with *HptMX6,* generating CWM345. The *tsa1::HptMX6* allele was then amplified from genomic DNA of CWM345 with oligonucleotides that flanked *TSA1* and used to generate *tsa1Δ* derivatives of the seven *CDC19* cysteine mutant strains via transformation.

### Protein extraction and immunoblotting

Protein extraction with trichloroacetic acid (TCA), protein detection by immunoblotting using a Li-Cor Odyssey infrared dye detection system, and protein quantitation were performed as previously described [20]. Rabbit antibodies specific to Tsa1 [61] were a gift of Shusuke Kuge. Antibody specific to Tsa1 hyperoxidized at C48 was provided by Ho Zoon Chae [32]. Rabbit anti-Pyk1 antibody [109] was a gift of Jeremy Thorner, and rabbit anti-Fba1 antibody [110] a gift of Magdalena Boguta. Rabbit anti-Ccs1 [111] was a gift of Valeria Culotta, rabbit anti-Hsp42 [112] was provided by Johannes Buchner, and rabbit anti-Pfk1/2 by Jurgen Heinisch [113]. Other antibodies were sourced from Invitrogen (rabbit Anti-MBP PA1-989), Abcam (mouse anti-Pgk1, 22C5D8, rabbit anti-GFP, ab290), Genscript (mouse anti-MBP, 00190-40), Nordic/MUBIO (rabbit anti-Hxk2, NE075/7S), Epigentek (rabbit anti-Pyc1, A53675), LS Bio (rabbit anti-Tdh1/2, LS-C147455) and Roche (mouse anti-GFP, 11814460001). IRDye^®^ 680LT dye-labeled anti-mouse antibody and IRDye^®^ 800CW dye-labeled anti-rabbit antibody were obtained from Li-Cor. Antibody specificity was verified by comparison of target proteins in wild-type and GFP-tagged or deletion mutant strains via immunoblotting (**Supplemental Fig. S7**).

### Purification of MBP fusions from yeast

Exponential phase yeast cultures (25-100 ml) were harvested by centrifugation and washed twice with 25 ml 1x PBS/1 mM EDTA. Unless otherwise indicated, cells were resuspended in 1x PBS/1 mM EDTA and treated with 10 mM NEM for 20 min at 30°C with shaking. Cells were collected by centrifugation and washed twice with 1 mM EDTA. Protein was extracted from cell pellets by addition of 0.5 ml extraction buffer containing 20 mM Tris–HCl pH 7.5, 200 mM NaCl, 5% Glycerol, 1% Triton X-100, 1x fungal protease inhibitor cocktail (Sigma P8215), 1 mM EDTA, 5 μM MG132, and 1 mM PMSF. Buffers also routinely contained 5 mM NEM. A 0.2 ml volume of glass beads was added and the tubes were vortexed for 10 min at 4°C. Tubes were centrifuged at 16,000 x g for 20 min at 4°C and the supernatant collected. The protein concentration of the supernatant was measured using Bradford reagent and adjusted to 2 mg/ml. MBP was purified from 0.5 - 1 mg of starting lysate protein using a spin column method. A chromatography column (Bio-Rad Bio-spin, 7326008) was supported by a 10 ml test tube support placed inside a 15 ml plastic tube. The column was loaded with 0.4 ml of amylose resin suspension (0.2 ml final resin volume) (NEB E8021S), 0.5 ml of extraction buffer was applied and the column centrifuged at 300 x g for 1 min at 4 °C. The wash was repeated two more times before the support tube was replaced with a 2 ml collection tube. Lysate protein (typically 0.5 ml of a 2 mg/ml solution) was loaded on the column and centrifuged as above. The column was washed four times with 0.5 ml ice cold extraction buffer by centrifugation, and bound proteins were eluted by addition of a 0.4 ml volume of extraction buffer + 10 mM maltose.

### Label free MS analysis of purified MBP fractions

For proteomic analysis samples of purified MBP and TRM were prepared as described above, except that cells were not subject to NEM treatment prior to lysis, and NEM in extraction buffer was replaced with 10 mM IAA. Label-free MS analysis of amylose column eluates was performed by the UW-Madison Mass Spectrometry/Proteomics Facility essentially as previously described [114]. All MS/MS samples were analyzed using Mascot (Matrix Science, London, UK; version 2.7.0). Mascot was set up to search the UniProtKB/SwissProt yeast database (SwissProt_YEAST_20201007) with general contaminants added (GPM cRAP database; cRAP_20190304), assuming the digestion enzyme trypsin. Mascot was searched with a fragment ion mass tolerance of 0.60 Da and a parent ion tolerance of 10.0 PPM. Carbamidomethylation of cysteine and j+138 of leucine/isoleucine indecision were specified in Mascot as fixed modifications. Deamidation of asparagine and glutamine and oxidation of methionine were specified in Mascot as variable modifications.

Scaffold (version 5.0.1, Proteome Software Inc., Portland, OR) was used to validate MS/MS based peptide and protein identifications. Peptide identifications were accepted if they could be established at greater than 91.0% probability to achieve an FDR less than 1.0% by the Scaffold Local FDR algorithm. Protein identifications were accepted if they could be established at greater than 5.0% probability to achieve an FDR less than 1.0% and contained at least 2 identified peptides. Protein probabilities were assigned by the Protein Prophet algorithm [115]. Proteins that contained similar peptides and could not be differentiated based on MS/MS analysis alone were grouped to satisfy the principles of parsimony. Proteins sharing significant peptide evidence were grouped into clusters.

### Quantitative proteomic analysis via stable isotope labeling by amino acids in cell culture (SILAC)

Protein accumulation in wild-type and *tsa1Δ* strains in zinc-deficient cells was compared using SILAC [116]. To enable metabolic labeling with lysine and arginine, a *lys2Δ0 arg4::KanMX4* yeast strain and an isogenic *tsa1Δ::LEU2* derivative were generated (see Experimental Procedures). Isotopically labeled *TSA1* wild-type cells were prepared by growth in LZM containing labeled arginine and lysine (^13^C6/^15^N4 arginine and ^13^C6/^15^N2 lysine, both 99%, Cambridge Isotope Laboratories Inc.) (“heavy” LZM). Lysine and arginine were supplied at a low concentration (50 mg/L) to minimize lysine conversion to other amino acids *in vivo* [117].

Uracil, histidine, and leucine (100 mg/L) were also added to medium to support auxotrophic requirements. The *tsa1Δ* mutant was supplied with the same medium containing arginine and lysine of natural isotopic distribution (“light” LZM). Initial overnight cultures of wild-type were started from single colonies in 5 ml heavy LZM, or for *tsa1Δ* in light LZM, both supplemented with 100 μM ZnCl_2_. Cultures were allowed to complete at least 7 generations of growth, harvested and washed with 1 mM EDTA to remove zinc, then used to inoculate four 20 ml aliquots of heavy (wild-type) or light (*tsa1Δ*) LZM supplemented with 0.7 μM ZnCl_2_, at an initial *A*_595_ of 0.01 (wild-type) or 0.04 (*tsa1Δ*) (zinc concentration in LZM was set at 0.7 μM to compensate for relatively high intracellular zinc stores resulting from growth in LZM + 100 Zn). Cells were then grown for another three generations, diluting with fresh medium as required to maintain cell density below 0.3 *A*_595_. A total of 6x10^7^ cells were collected and resuspended in soluble protein buffer (50 mM Tris-Cl pH 8.5, 500 mM NaCl, 10 mM EDTA, 1x fungal protease inhibitor cocktail [Sigma P8215], 1 mM EDTA, 5 μM MG132, and 1 mM PMSF). Samples were frozen with liquid N_2_ and disrupted using a mixer mill (Retsch) [9]. Protein was precipitated by addition of TCA to 10%, washed twice with acetone, and redissolved in denaturing buffer (100 mM Tris-Cl pH 8.5, 2% SDS, 150 mM NaCl, 1 mM EDTA). Samples were centrifuged at 14,000 x*g* for 10 min to remove insoluble material and the concentration measured with a DC assay kit (#5000111, Bio-Rad). Two hundred μg of protein from each wild-type and *tsa1Δ* replicate were combined to give four replicate mixtures. A 1.5x volume of 10 M urea was added to each sample, and SDS was removed by passage through a 0.5 ml detergent removal column (#87777, Pierce) equilibrated with Urea buffer (100 mM Tris pH 8.5. 150 mM NaCl, 6M urea, 1 mM EDTA). The protein was precipitated again with TCA and pellets washed with acetone prior to MS analysis.

For Tandem LC-MS/MS analysis, protein samples resuspended in digestion buffer (100 mM Tris pH 8.5, 8M urea) were reduced and alkylated by sequential incubations with 5 mM TCEP-HCl for 20 min at room temperature followed by treatment with 10 mM iodoacetamide for 20 min at room temperature in the dark. Proteins were then digested to peptides by incubating with Lys-C at an enzyme-to-substrate ratio of 1:100 for 4 hours at 37 °C, followed by dilution to 2 M urea using 100 mM Tris, pH 8.5 and incubation with trypsin at an enzyme-to-substrate ratio of 1:50 for 12 additional hours at 37 °C. The digest was stopped by the addition of formic acid at a final concentration of 5%, and desalted using C18 tips (Pierce) according to the manufacturer’s instructions. For LC–MS/MS analysis, digested peptides were separated online using C18 reversed phase chromatography with an EASY-nLC 1000 ultra-high-pressure liquid chromatography (UHPLC) system (Thermo Fisher) coupled to a 75 μM inner diameter fused silica capillary column with a 5 μM integrated emitter and packed in-house with 15 cm of Luna C18(2) 3 μm reversed-phase particles [118]. MS/MS spectra were collected in data-dependent mode on a Q Exactive mass spectrometer (Thermo Fisher) [119]. Data analysis was performed with the ProLuCID and DTASelect2 algorithms implemented in the Integrated Proteomics Pipeline - IP2 (Integrated Proteomics Applications, San Diego, CA) considering both light and heavy versions of each peptide [120, 121]. Peptide spectrum matches were filtered with a false positive rate of <5% as estimated by a decoy database strategy [122]. SILAC ratios were calculated by the Census algorithm [123].

### Gene Ontology analysis

FunSpec (funspec.med.utoronto.ca) was used to search the Tsa1 interactome for significantly overrepresented GO molecular function, biological process, and cellular component terms, as well as MIPS functional classifications. Searches were performed with a *p*-value cutoff of 0.01 and Bonferroni correction applied.

### Data availability

All data generated in this study are available by request to CWM. MS proteomic data will be deposited in the ProteomeXchange through PRIDE.

## ACKNOWLEDGEMENTS

The authors wish to thank Amanda Bird for plasmid pYEF2-myc-Zap1^642-880^ and the ZAP1-13xmyc strain, Andrew Herbig for plasmid pAFH13, and Audrey Gasch for her gift of multiple GFP-tagged yeast strains. We also thank Shusuke Kuge, Ho Zoon Chae, Jeremy Thorner, Magdalena Boguta ,Valeria Culotta, Johannes Buchner and Jurgen Heinisch for their generous gifts of antibodies and plasmids used in this study, and Greg Sabat and Greg Barrett-Wilt at the UW-Madison Mass Spectrometry Facility for their invaluable assistance with sample analysis and data processing.

## FUNDING AND ADDITIONAL INFORMATION

This work was supported by National Institutes of Health grant R01-GM056285 (DJE) and R35-GM153408 (JAW). The content is solely the responsibility of the authors and does not necessarily represent the official views of the National Institutes of Health.

## SUPPLEMENTAL TABLES

Table S1 Complete interactome dataset.

Table S2 Targets enriched >3-fold ZnR and ZnD.

Table S3 Biogrid comparison.

Table S4 GO analysis.

Table S5 Tsa1 targets in amino acid pathways.

**Table S6.**
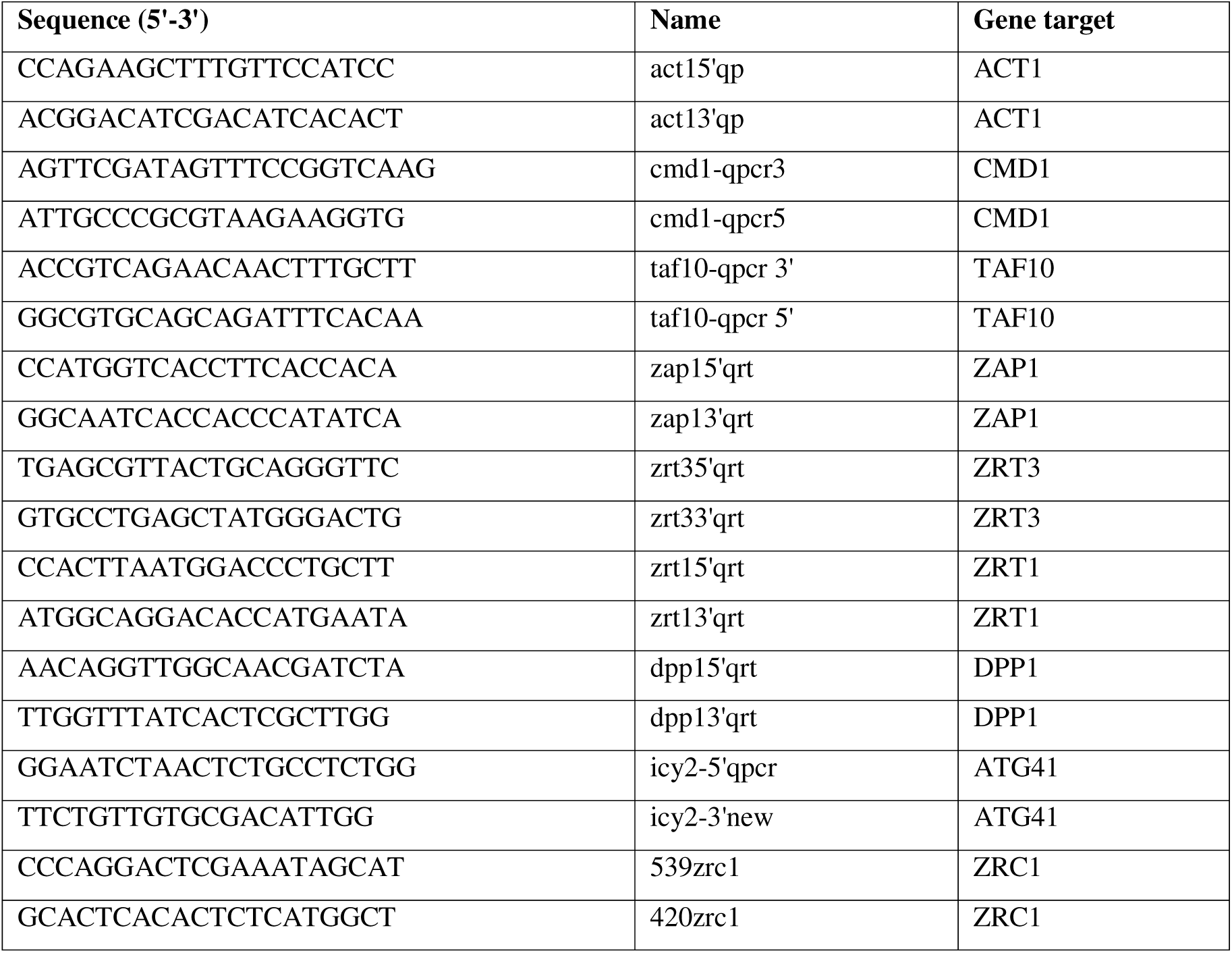
Oligonucleotides.

**Table S7.**
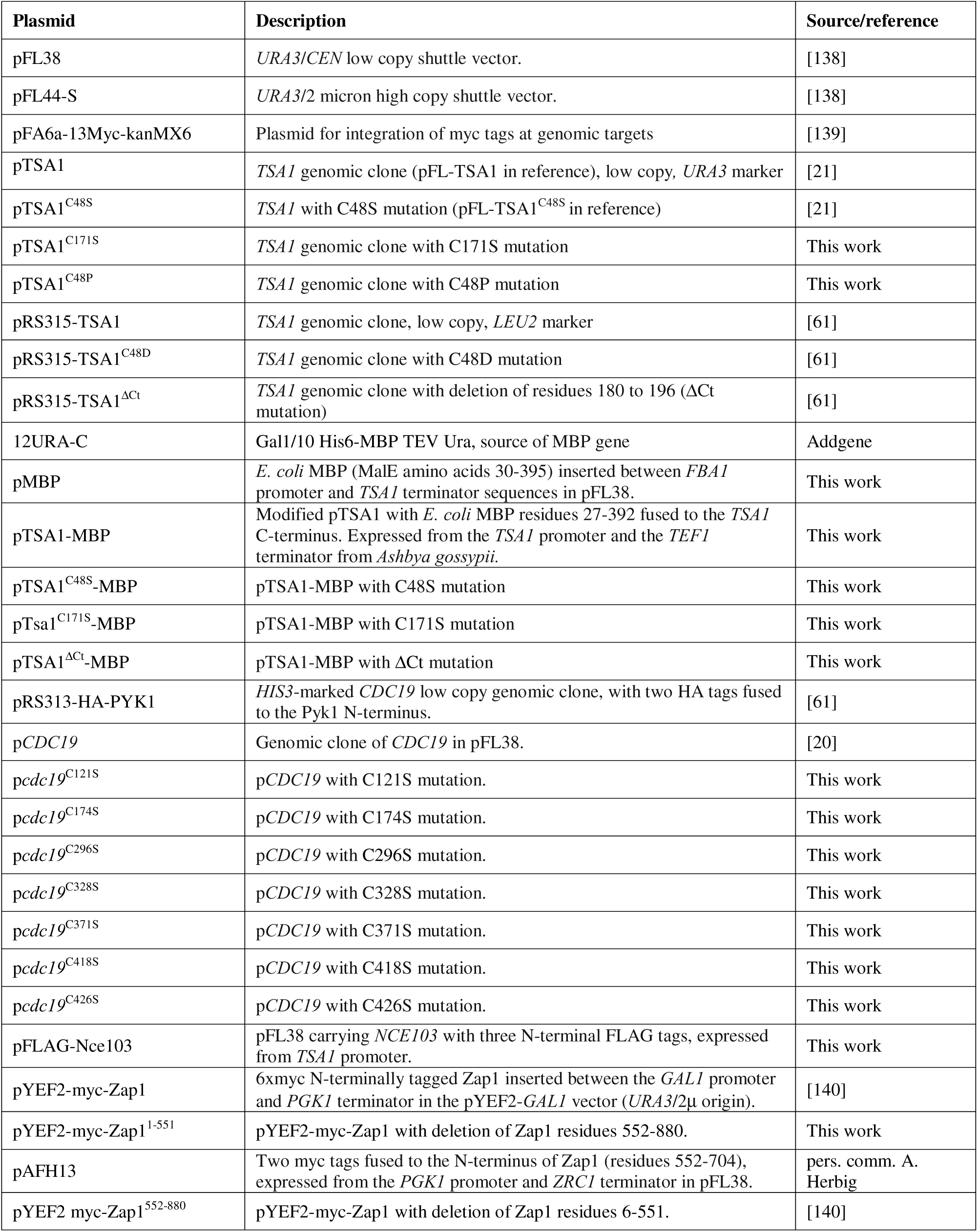
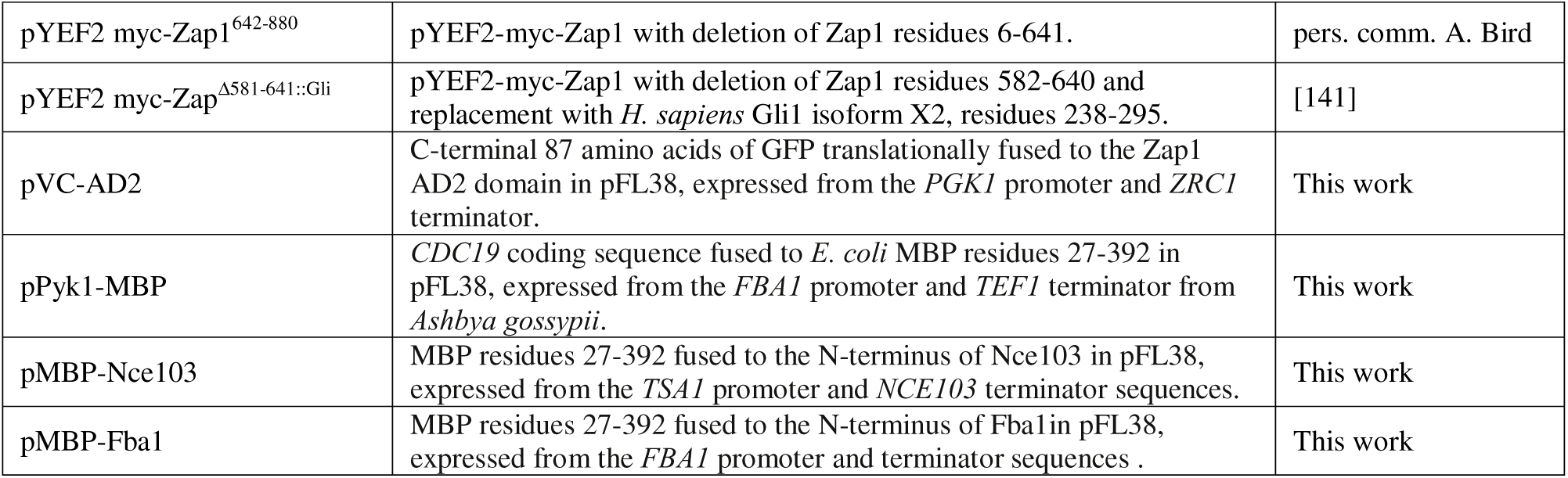
Plasmids.

**Table S8.** Yeast strains.

| Strain | Relevant genotype | Full genotype | Source/reference |
| --- | --- | --- | --- |
| BY4741 | WT | *MATa his3Δ0 leu2Δ0 met15Δ0 ura3Δ0* | [106] |
| BY4742 | WT | *MATα his3Δ0 leu2Δ0 lys2Δ0 ura3Δ0* | [106] |
| BY4743 | WT | *MATa/MATα his3Δ1/his3Δ1 leu2Δ0/leu2Δ0 met15Δ0/MET15 lys2Δ0/LYS2 ura3Δ0/ura3Δ0* | [106] |
| BY4742/*tsa1Δ* | *tsa1Δ* | *MATα tsa1::KanMX4 his3Δ0 leu2Δ0 lys2Δ0 ura3Δ0* | [106] |
| ABY64 | *ADH4-3xHA::TRP1* | *MATa his3Δ1 leu2Δ0 met15Δ0 ura3Δ0 trp1::KanMX4* | [142] |
| CWM231 | *TEF2pr-GFP* | *MATa his3::TEF2pr-GFP-NatMX4 his3Δ0 leu2Δ0 met15Δ0 ura3Δ0* | This work |
| BY4742/*arg4Δ* | *arg4Δ lys2Δ0* | *MATα arg4::KanMX4 his3Δ0 leu2Δ0 lys2Δ0 ura3Δ0* | [106] |
| CWM254 | *arg4Δ lys2Δ0 tsa1Δ* | *MATα arg4::KanMX4 tsa1::LEU2 his3Δ0 leu2Δ0 lys2Δ0 ura3Δ0* | This work |
| CWM345 | *tsa1::HptMX6* | *MATα tsa1::HphMX6 his3Δ0 leu2Δ0 lys2Δ0 ura3Δ0* | This work |
| CWM347 | *ZAP1-13xmyc tsa1Δ* | *MATa ZAP1-13xmyc::KanMX6 tsa1::LEU2 his3Δ0 leu2Δ0 met15Δ0 ura3Δ0* | This work |
| CWM355 | *ARO4-GFP tsa1Δ* | *MATα ARO4-GFP tsa1::KanMX4 his3Δ0 leu2Δ0 lys2Δ0 ura3Δ0* | This work |
| CWM357 | *cdc19<sup>C174S</sup> tsa1Δ* | *MATα cdc19<sup>C174S</sup> tsa1::HphMX6 his3Δ0 leu2Δ0 lys2Δ0 ura3Δ0* | this work |
| CWM359 | *cdc19<sup>C296S</sup> tsa1Δ* | *MATα cdc19<sup>C296S</sup> tsa1::HphMX6 his3Δ0 leu2Δ0 lys2Δ0 ura3Δ0* | this work |
| CWM361 | *cdc19<sup>C328S</sup> tsa1Δ* | *MATα cdc19<sup>C328S</sup> tsa1::HphMX6 his3Δ0 leu2Δ0 lys2Δ0 ura3Δ0* | this work |
| CWM363 | *cdc19<sup>C371S</sup> tsa1Δ* | *MATα cdc19<sup>C371S</sup> tsa1::HphMX6 his3Δ0 leu2Δ0 lys2Δ0 ura3Δ0* | this work |
| CWM365 | *cdc19<sup>C418S</sup> tsa1Δ* | *MATα cdc19<sup>C418S</sup> tsa1::HphMX6 his3Δ0 leu2Δ0 lys2Δ0 ura3Δ0* | this work |
| CWM367 | *cdc19<sup>C426S</sup> tsa1Δ* | *MATα cdc19<sup>C426S</sup> tsa1::HphMX6 his3Δ0 leu2Δ0 lys2Δ0 ura3Δ0* | this work |
| CWM384 | *ADH4-3xHA::TRP1 tsa1Δ* | *MATa tsa1::LEU2 his3Δ1 leu2Δ0 met15Δ0 ura3Δ0 trp1::KanMX4* | this work |
| CWM442 | *cdc19<sup>C121S</sup> tsa1Δ* | *MATα cdc19<sup>C121S</sup> tsa1::HphMX6 his3Δ0 leu2Δ0 lys2Δ0 ura3Δ0* | this work |
| CWM443 | *cdc19<sup>C328S</sup> tsa1Δ* | *MATα cdc19<sup>C328S</sup> tsa1::HphMX6 his3Δ0 leu2Δ0 lys2Δ0 ura3Δ0* | this work |
| BY4742/ARO4-GFP | *ARO4-GFP* | *MATα ARO4-GFP::His3MX6 his3Δ0 leu2Δ0 lys2Δ0 ura3Δ0* | [107] |
| ZAP1-13xmyc | *ZAP1-13xmyc* | *MATa ZAP1-13xmyc::KanMX6 his3Δ0 leu2Δ0 met15Δ0 ura3Δ0* | pers. comm. A. Bird |
| BY4742/TSA1-GFP | *TSA1-GFP* | *MATα TSA1-GFP::His3MX6 his3Δ0 leu2Δ0 lys2Δ0 ura3Δ0* | [107] |
| BY4742/CCS1-GFP | *CCS1-GFP* | *MATα CCS1-GFP::His3MX6 his3Δ0 leu2Δ0 lys2Δ0 ura3Δ0* | [107] |
| BY4742/HSP42-GFP | *HSP42-GFP* | *MATα HSP42-GFP::His3MX6 his3Δ0 leu2Δ0 lys2Δ0 ura3Δ0* | [107] |
| BY4742/PFK1-GFP | *PFK1-GFP* | *MATα PFK1-GFP::His3MX6 his3Δ0 leu2Δ0 lys2Δ0 ura3Δ0* | [107] |
| BY4742/HXK1-GFP | *HXK1-GFP* | *MATα HXK1-GFP::His3MX6 his3Δ0 leu2Δ0 lys2Δ0 ura3Δ0* | [107] |
| BY4742/PYC1-GFP | *PYC1-GFP* | *MATα PYC1-GFP::His3MX6 his3Δ0 leu2Δ0 lys2Δ0 ura3Δ0* | [107] |

## SUPPLEMENTAL FIGURES

**Supplemental Figure S1.**
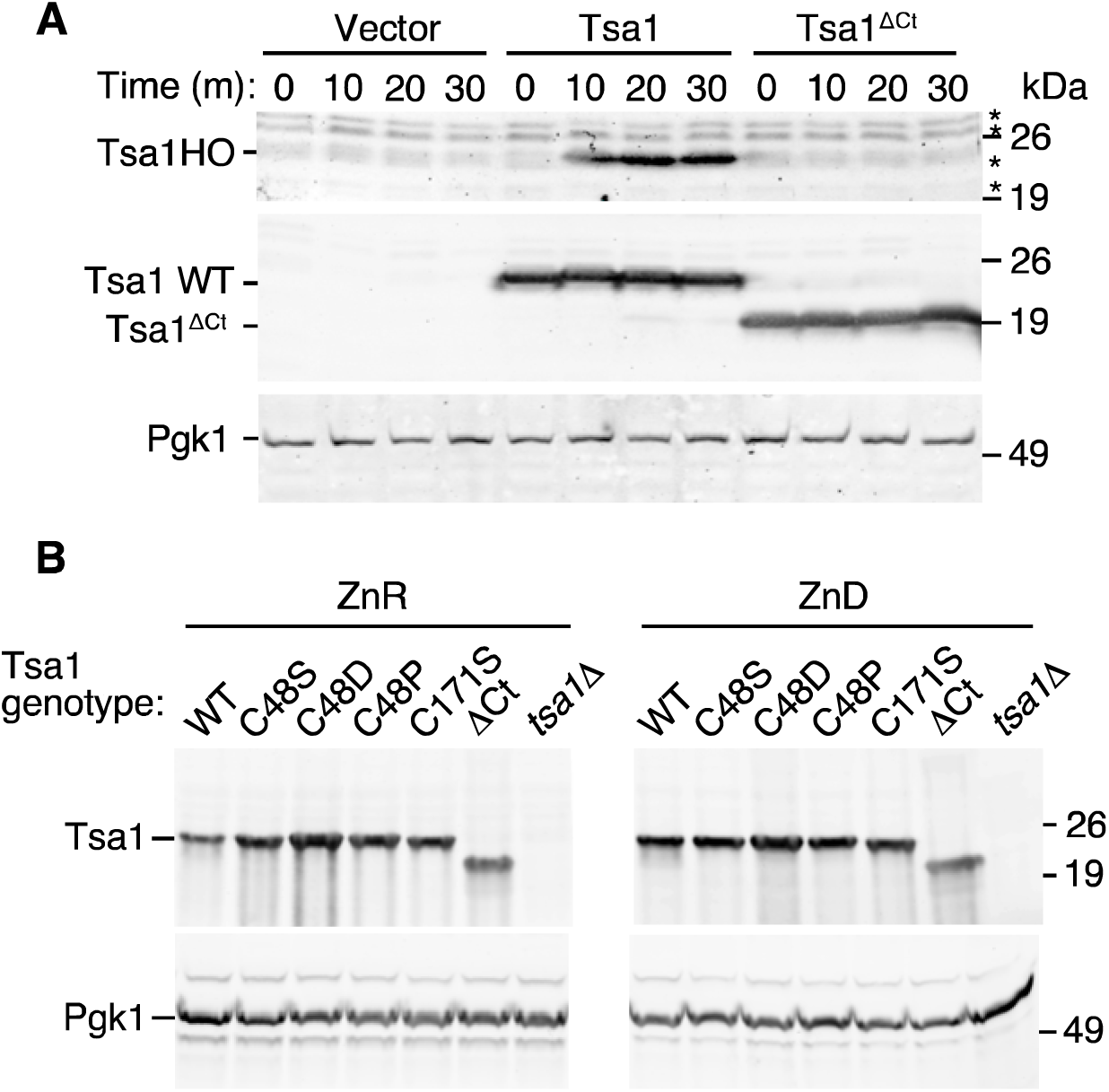
Verification of Tsa1 mutant phenotypes. A) The Tsa1^ΔCt^ mutation prevents hyperoxidation. BY4742 *tsa1Δ* was transformed with pRS315 (vector control), pRS315-Tsa1 (WT), or pRS315-Tsa1^ΔCt^ plasmids. Strains were grown to log phase in SD medium and treated with 5 mM H_2_O_2_ for the indicated times. Protein was extracted using TCA and subjected to immunoblot analysis to detect hyperoxidized Tsa1 (*top*) or total Tsa1 (*middle*). Asterisks indicate nonspecific background bands. B) Accumulation of mutant Tsa1 proteins. BY4742 *tsa1Δ* was transformed with vector alone (pFL38, *tsa1Δ*), wild-type Tsa1 (pTSA1), or the indicated *TSA1* mutant alleles (plasmids pTSA1^C48S^, pRS315-TSA1^C48D^, pTSA1^C48P^, pTSA1^C171S^, or pRS315-TSA1^ΔCt^). Strains were grown in zinc-replete (ZnR, LZM + 100 μM ZnCl_2_) or deficient (ZnD, LZM + 1 μM ZnCl_2_) conditions with appropriate selection (-uracil or - leucine), and total protein extracts subjected to immunoblotting with anti-Tsa1. In A and B, Pgk1 was detected as a loading control.

**Supplemental Figure S2.**
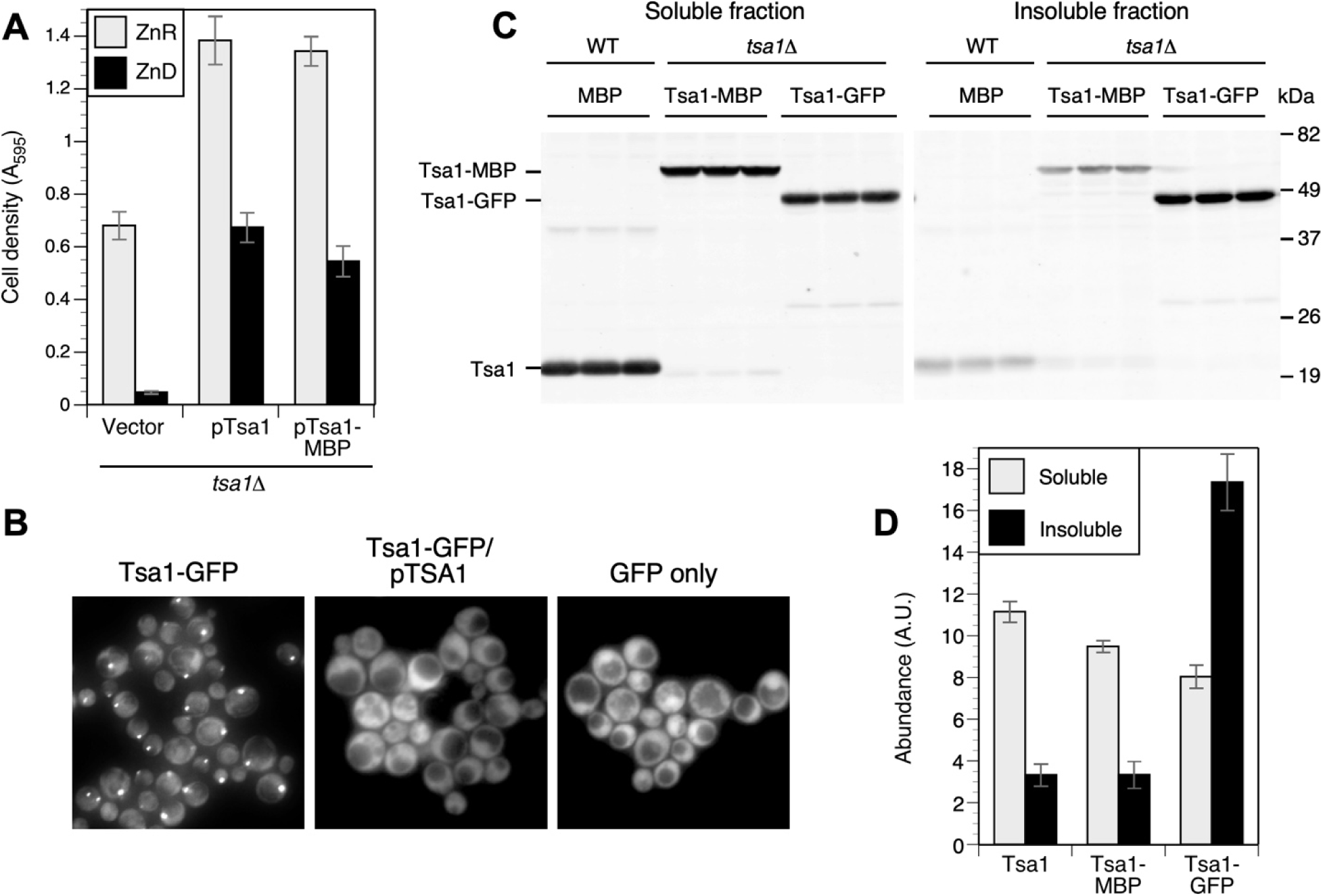
Verification of Tsa1-MBP function and stability. A) The Tsa1-MBP fusion complemented the *tsa1Δ* growth defect in low zinc. A *tsa1Δ* mutant (BY4742/*tsa1*) was transformed with the indicated plasmids and the resulting strains were used to inoculate zinc-replete (LZM+100 μM ZnCl_2_) or zinc-deficient (LZM + 1 μM ZnCl_2_) medium to an initial A_595_ of 0.01. Cell density was measured after 16 h growth in ZnR or 2 days in ZnD medium. B) Tsa1-GFP aggregated in ZnD cells. Aliquots of ZnD medium (LZM + 1 μM ZnCl_2_) were inoculated with BY4742 expressing Tsa1 tagged with GFP at the endogenous locus (BY4742/Tsa1-GFP), the same strain transformed with a plasmid expressing untagged Tsa1 (Tsa1-GFP/pTsa1), or a strain expressing GFP alone from the *TEF2* promoter (CWM231). Cells were grown to log phase and GFP imaged with a wide-field fluorescence microscope. Images are representative of three independent experiments. C) Tsa1-GFP was less soluble than untagged Tsa1 in ZnD cells, but Tsa1-MBP solubility was little affected. Wild-type BY4742 transformed with the pMBP plasmid (MBP), or congenic *tsa1Δ* mutant strains transformed with pTsa1-MBP or pTsa1-GFP plasmids were grown in LZM + 1 μM ZnCl_2_, and detergent-resistant insoluble fractions isolated as described in Experimental Procedures. Immunoblots of yeast lysates (*left* panel) or detergent-resistant fractions (*right* panel) were probed with antibody to Tsa1. Three experimental replicates for each genotype are shown. D) Relative abundance in arbitrary units (A. U.) of Tsa1 and tagged derivatives in ZnD fractions quantified from the immunoblot in panel C. Values are the averages of three replicates and error bars indicate ± 1 S.D.

**Supplemental Figure S3.**
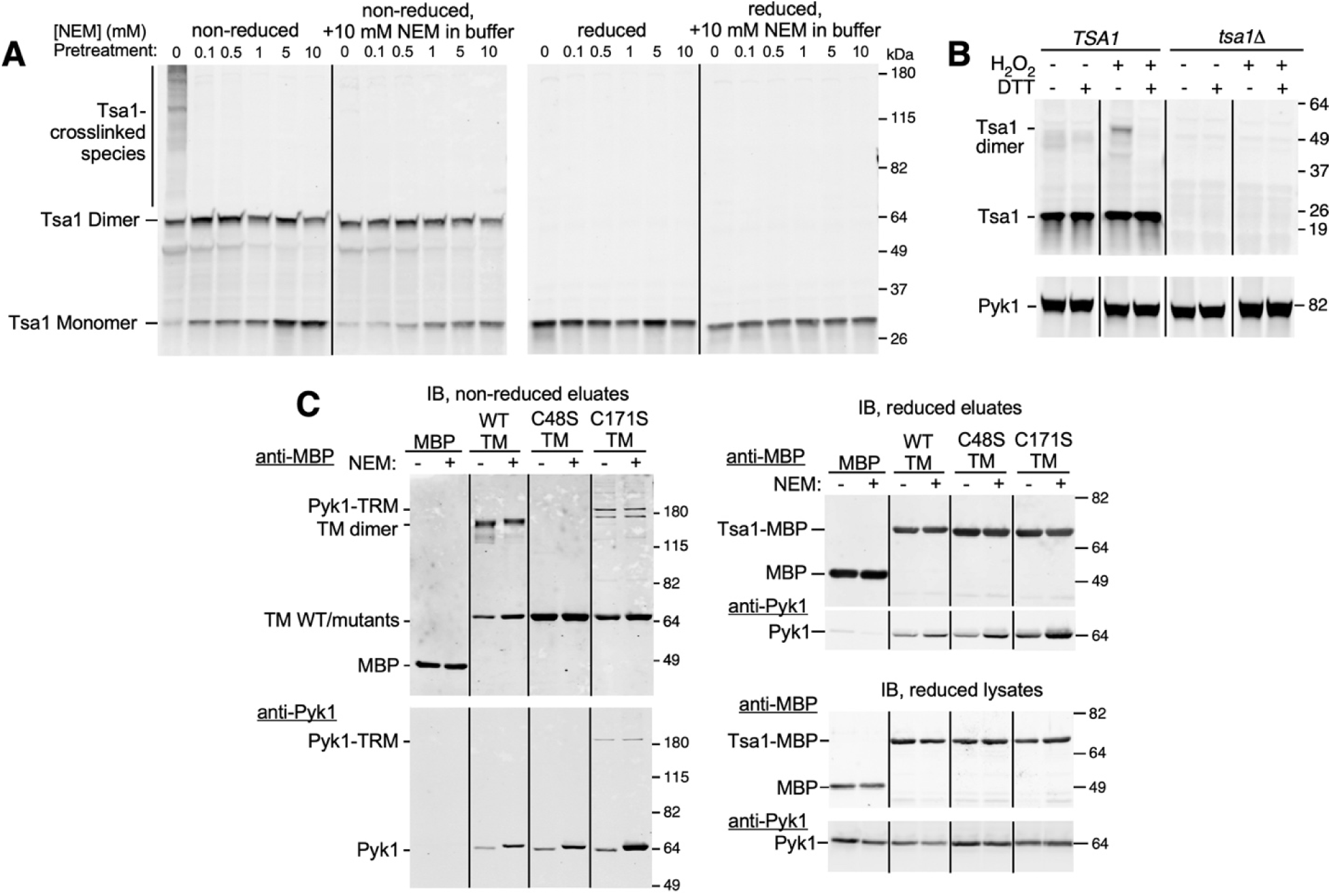
Effect of NEM on Tsa1 oxidation and interaction with Pyk1. A) Wild-type (BY4742) ZnR cells were treated with the indicated concentration of NEM for 15 min and lysates prepared with extraction buffer that contained no or 10 mM NEM as indicated. Non-reduced (*left* panel) or reduced (β−mercaptoethanol-treated, *right* panel) samples were analyzed by immunoblotting and Tsa1 detected with anti-Tsa1. B) Zinc-replete wild-type (BY4742) or congenic *tsa1Δ* mutant cells were treated with 0.5 mM H_2_O_2_ for 10 min where indicated, a concentration that minimized hyperoxidation. Cells were then treated with TCA and protein extracted to preserve redox state. Equal amounts of protein were reduced with DTT where indicated, fractionated in a 4-12% gradient SDS-PAGE gel, and analyzed by immunoblotting with anti-Tsa1 and anti-Pyk1 antibodies. C) WT cells (BY4742) expressing MBP alone, or *tsa1Δ* mutant cells expressing wild-type or the indicated mutant alleles of Tsa1-MBP (C48S or C171S) were grown in zinc-replete medium, and treated with 5 mM NEM where indicated. Protein was extracted for MBP purification, and eluate or lysate samples were fractionated on a 10% SDS-PAGE gel and analyzed by immunoblotting to detect MBP and Pyk1. Note that the difference in apparent molecular mass of Pyk1 in panels B and C is likely a consequence of the difference in SDS-PAGE systems used.

**Supplemental Figure S4.**
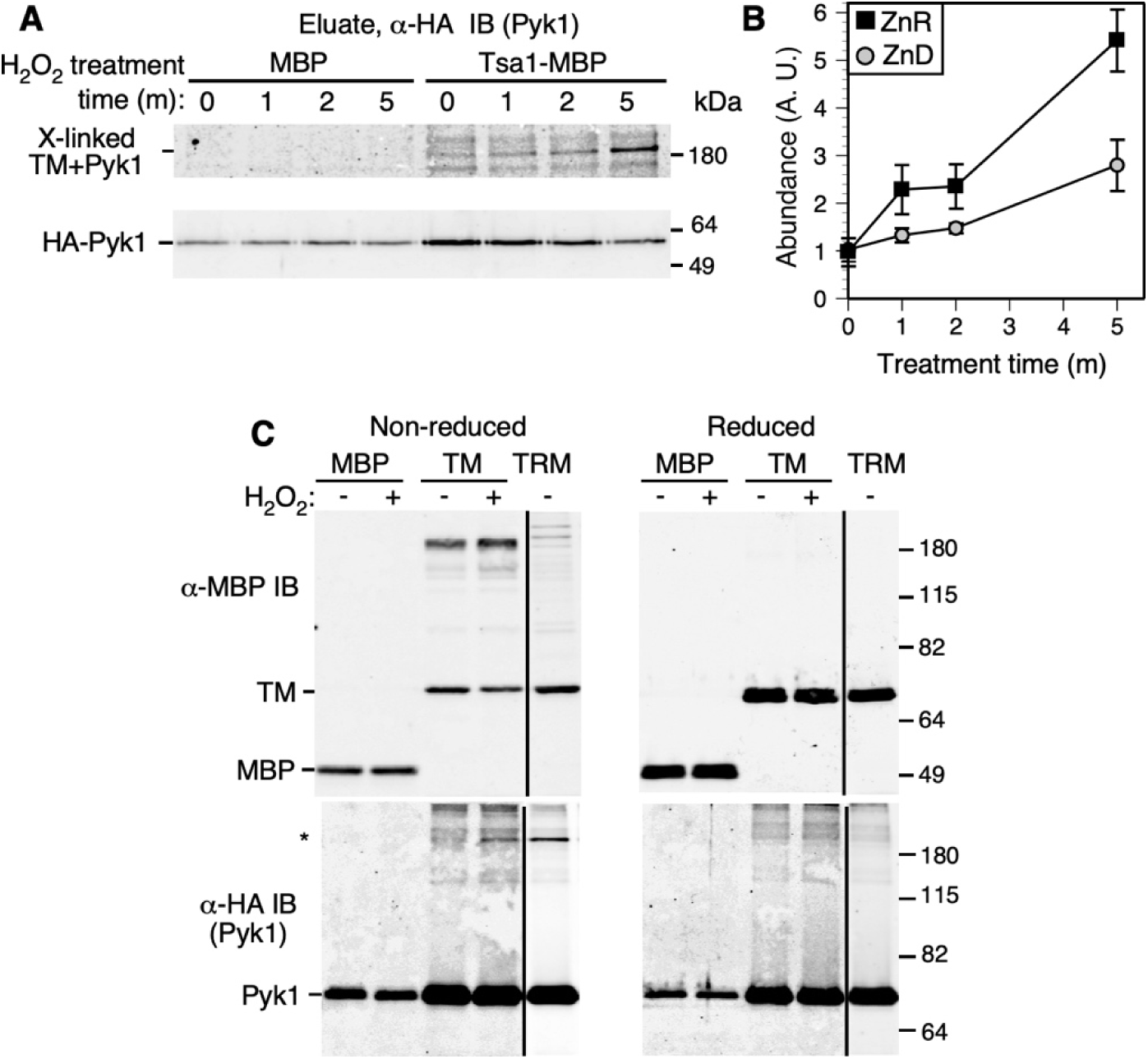
Wild-type Tsa1 formed a mixed disulfide with Pyk1 in response to peroxide treatment *in vivo*. A) Wild-type cells (BY4742) expressing HA-tagged Pyk1 and MBP, or *tsa1Δ* cells expressing HA-tagged Pyk1 and wild-type Tsa1-MBP (plasmids pTSA1-MBP and pRS313-HA-PYK1) were grown to log phase in ZnR medium (LZM+100 μM ZnCl_2_) and either not treated (0 time), or treated with 100 μM H_2_O_2_ for the indicated times. Reaction of peroxide with cysteine thiols was terminated by addition of 10 mM NEM, native cell lysates prepared, and MBP purified. Eluate fractions were subject to immunoblotting to detect HA-tagged Pyk1. Positions of putative mixed disulfide of HA-Pyk1 and Tsa1-MBP as determined by immunoblot overlay (non-reduced samples, *upper* panel) and HA-Pyk1 alone (reduced samples, *lower* panel) are indicated. B) Abundance of the mixed disulfide species formed after H_2_O_2_ treatment of ZnR and ZnD cells was quantified from immunoblots. Data points are the average of three experimental replicates, including that shown in panel A (A.U. = arbitrary units). Error bars indicate ± 1 S.D. C) The H_2_O_2_-induced HA-Pyk1 band is reduction sensitive and migrates identically to the Pyk1-TRM mixed disulfide in SDS-PAGE. The strains described in panel A, as well as a *tsa1* strain expressing HA-Pyk1 and TRM were treated for 5 min with 100 μM H_2_O_2_ (or untreated as indicated) before oxidation was terminated by addition of 10 mM NEM. Immunoblots of purified MBP fractions were performed as described for panel A. Protein samples were untreated (*left* panel) or reduced by treatment with β-mercaptoethanol (*right* panel) prior to SDS-PAGE. Position of the Pyk1+Tsa1-MBP (or TRM) mixed disulfide is indicated by an asterisk. Note the background “smear” of reduction-resistant Pyk1 in this region of the gel likely represents SDS-resistant aggregates.

**Supplemental Figure S5.**
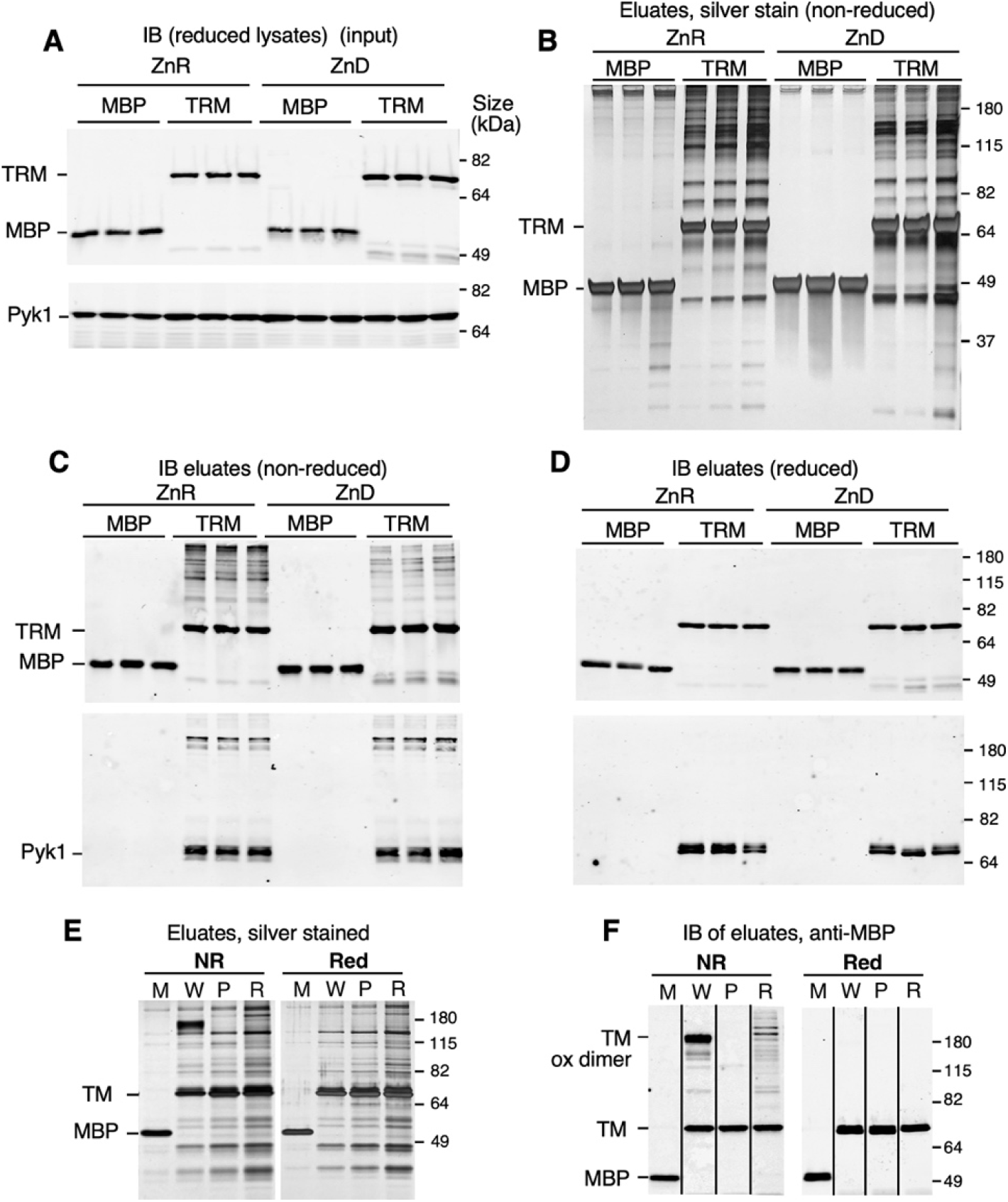
Analysis of samples for mass spectrometry of the Tsa1 interactome. A) Abundance of Tsa1^C171S^-MBP (TRM) and Pyk1 in cell lysates. Wild-type cells (BY4742) expressing MBP alone, or an isogenic *tsa1Δ* mutant expressing TRM were grown to log phase in ZnR or ZnD medium and native protein extracts prepared. Immunoblot shows the level of MBP, TRM, and Pyk1 proteins in triplicate replicates of reduced lysate samples. B) Composition of amylose column eluates. Equal amounts of input protein (2 mg) were loaded on amylose resin columns for purification fo MBP or TRM. Equal volumes of non-reduced column eluates were then fractionated by SDS-PAGE (10%) and silver stained. C, D) Immunoblots of non-reduced (C) or reduced (D) eluate samples were probed with antibodies against MBP (*upper* panels) or Pyk1 (*lower* panels). E, F) Effect of sample reduction on the composition of Tsa1-MBP eluates. MBP or Tsa1-MBP were purified from ZnR cells expressing wild-type Tsa1-MBP of the indicated mutant version (M = MBP alone, W = wild-type Tsa1-MBP, P = C48S, R = C171S). Cell lysis buffer contained 10 mM IAA to alkylate cysteines during extraction. Sample aliquots were reduced with 10 mM DTT (or not reduced, NR) prior to SDS-PAGE, and protein visualized by silver staining (E), or gel immunoblots probed with anti-MBP (F).

**Supplemental Figure S6.**
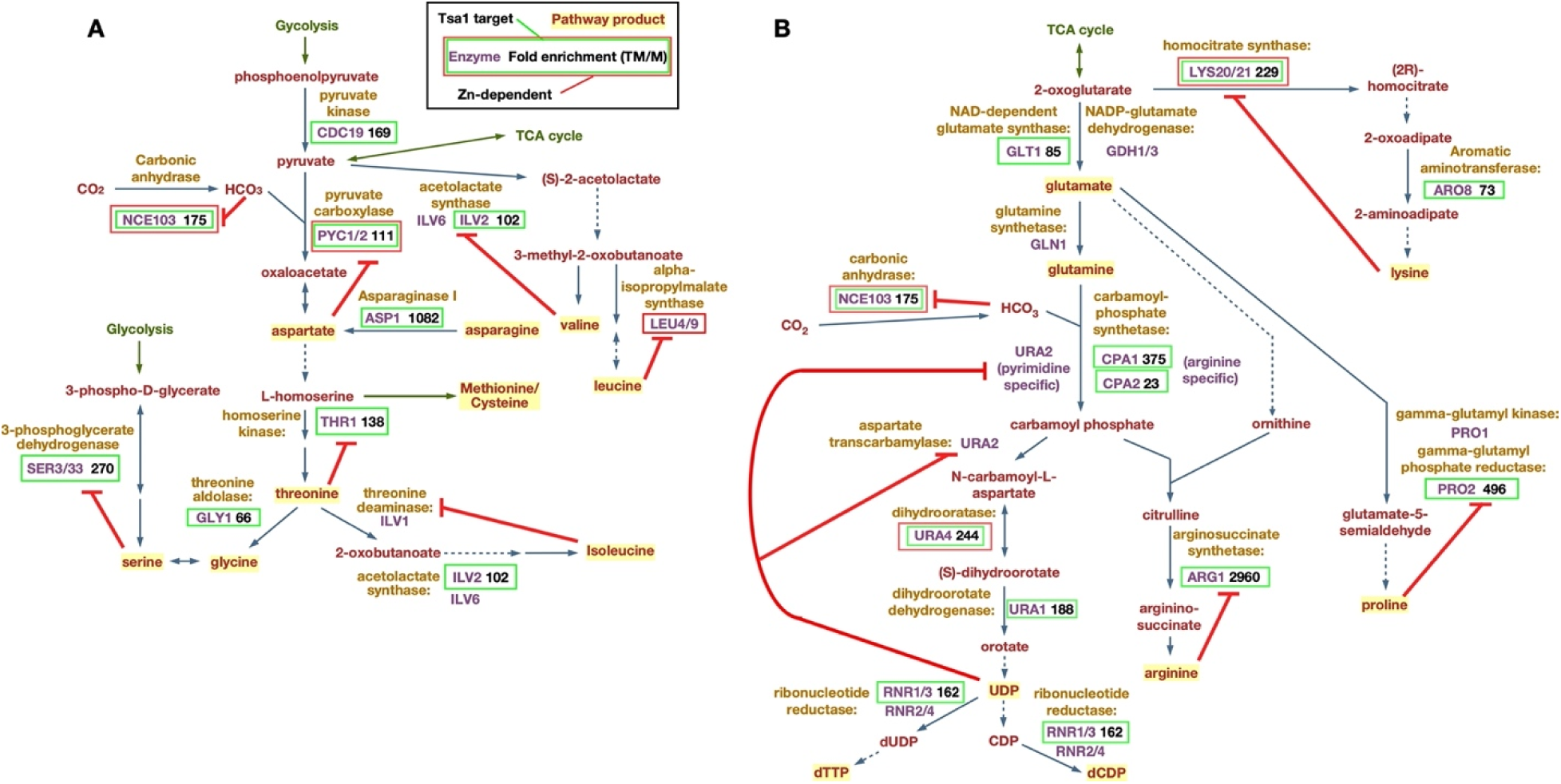
**Tsa1 targets in amino acid and pyrimidine synthesis pathways**. A) Pathways for synthesis of the aspartate family of amino acids. B) Pathways for synthesis of lysine, glutamate, glutamine, proline, arginine and pyrimidines. As indicated by legend (A, *top*), Tsa1 interactome components are outlined in *green*, and zinc-dependent enzymes are outlined in *red*. For each target, fold enrichment in TRM vs MBP eluate fractions is indicated (values are the average of ZnR and ZnD samples). Red lines indicate enzyme inhibition by pathway products.

**Supplemental Figure S7.**
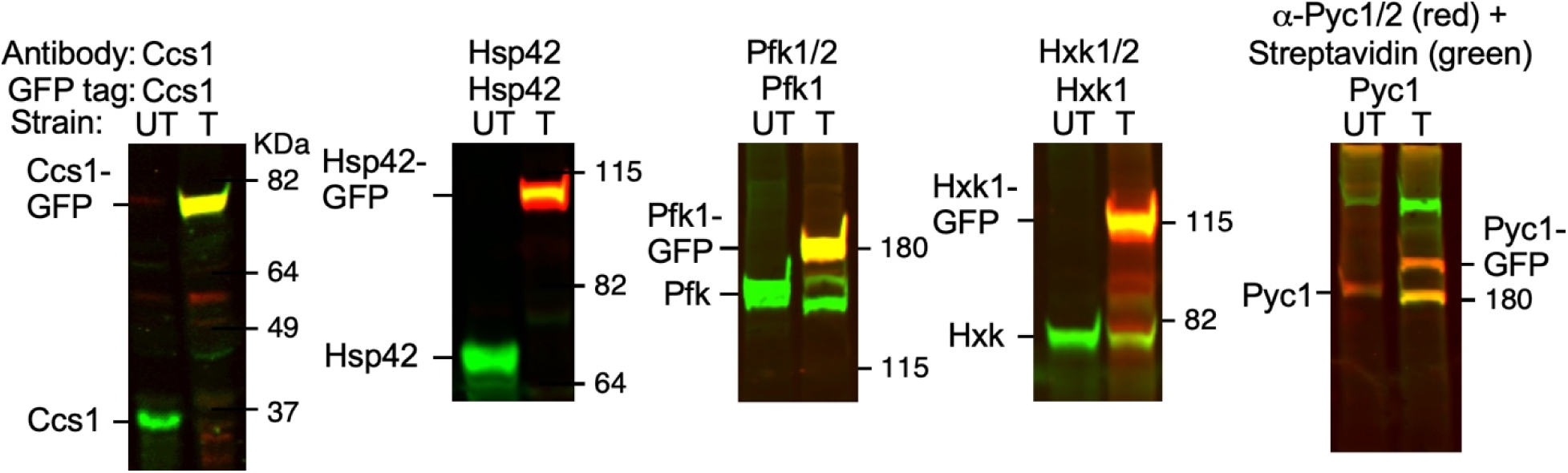
Verification of antibody specificity. Specificity of antibodies was verified by detection of novel bands reacting with both the specific antibody (*green* channel) and anti-GFP (*red* channel) in immunoblot experiments. Cells of GFP-tagged (T) or untagged (UT, BY4742) were grown in SD medium and protein extracted using TCA. Position of molecular mass markers is shown on the right of each panel. The antibody to pyruvate carboxylase (Pyc1/2) detected another unrelated band and the correct Pyc1/2-containing bands were identified via the binding of dye-labelled streptavidin to the Pyc1/2 biotin moiety.

**Supplemental Figure S8.**
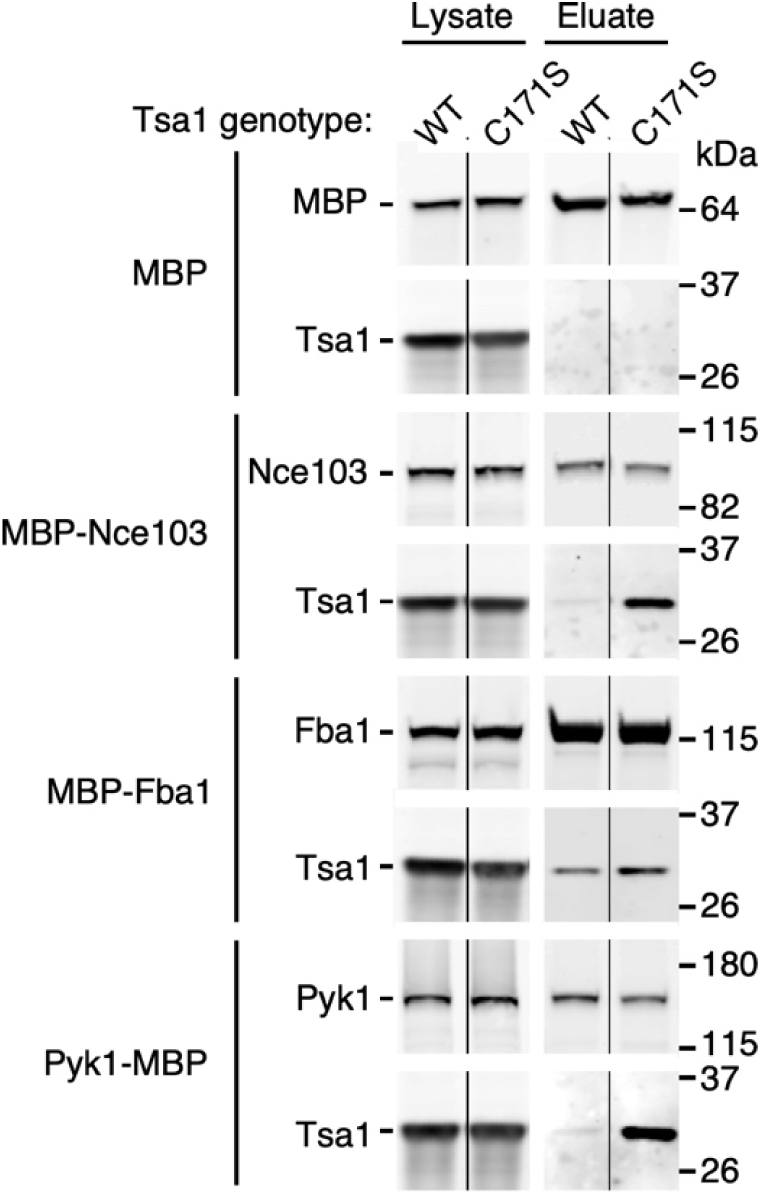
Tsa1 interacts with MBP-tagged targets. Cultures of BY4742/*tsa1* cells were transformed with plasmids expressing MBP alone, or MBP-tagged Fba1, Nce103 and Pyk1, together with myc-tagged wild type Tsa1 or the C171S mutant (pmyc-Tsa1 and pmyc-Tsa1^C171S^ plasmids respectively). Strains were grown to log phase in zinc-replete medium (LZM + 100 μM ZnCl_2_), MBP was purified, and the lysate and eluate fractions subjected to SDS-PAGE and immunoblotting to detect MBP-tagged proteins and myc-tagged Tsa1. Input levels of protein are also shown (lysates). Data shown are representative of two independent experiments.

**Supplemental Figure S9.**
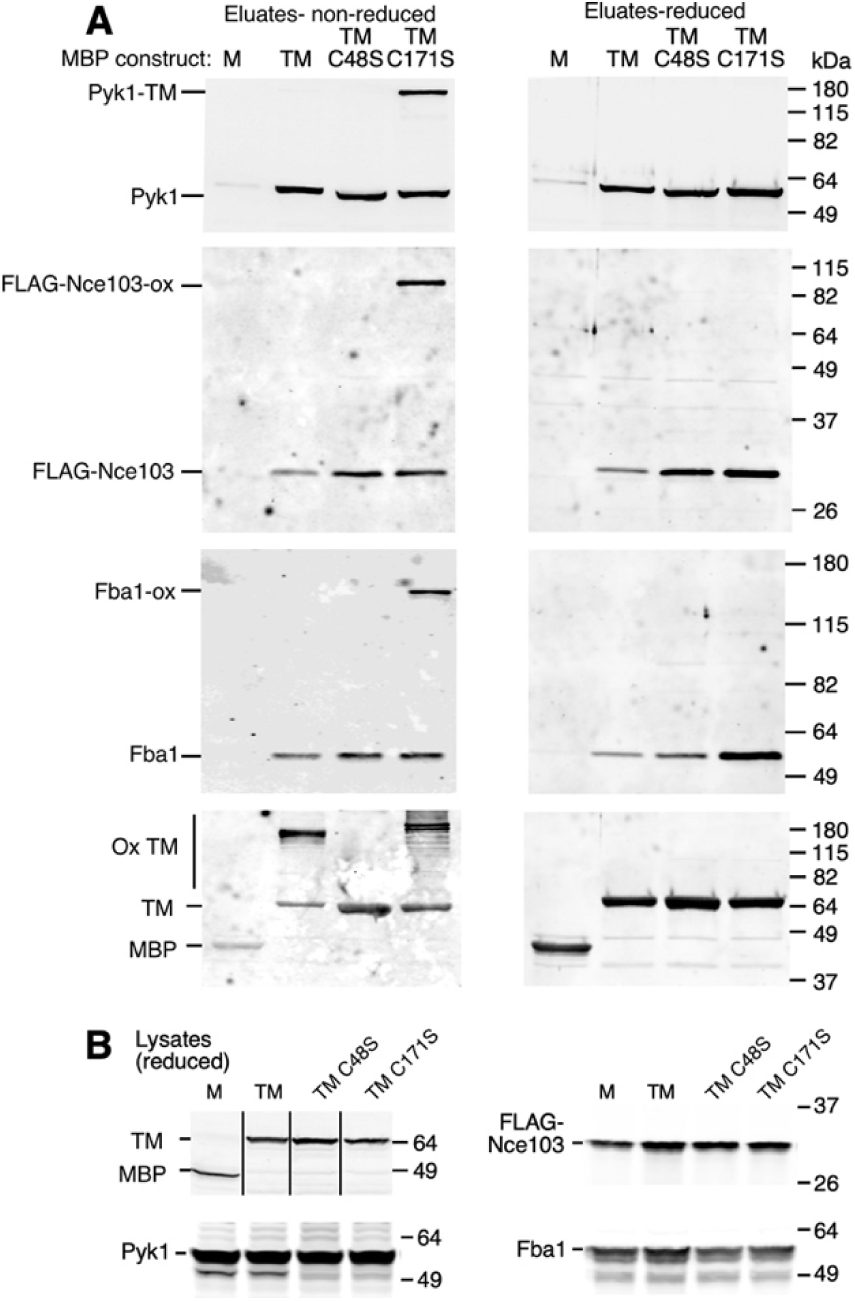
Detection of apparent mixed disulfides of Tsa1^C171S^-MBP and target proteins. A) Wild-type BY4742 was transformed with pFLAG-NCE103 and MBP alone (M), and a congenic *tsa1Δ* mutant strain was transformed with pFLAG-Nce103 and pTsa1-MBP (TM), pTsa1^C48S^-MBP (TMC48S) or pTsa1^C171S^-MBP (TMC171S). The resulting four strains were grown to log phase in LZM + 100 μM ZnCl_2_ and protein extracted. Cells were not treated with NEM prior to extraction, but 5 mM NEM was included in the extraction buffer. Equal volumes of non-reduced (A, *left* panel) or reduced (A, *right* panel) eluates were fractionated on 10% SDS-PAGE and immunoblots probed with antibodies for the indicated proteins. B) Protein abundance in reduced lysate samples was detected by immunoblotting as described for panel A. Lower molecular mass bands are minor degradation products. Positions of molecular mass standards are indicated on the right. Vertical lines indicate removal of superfluous lanes.

**Supplemental Figure S10.**
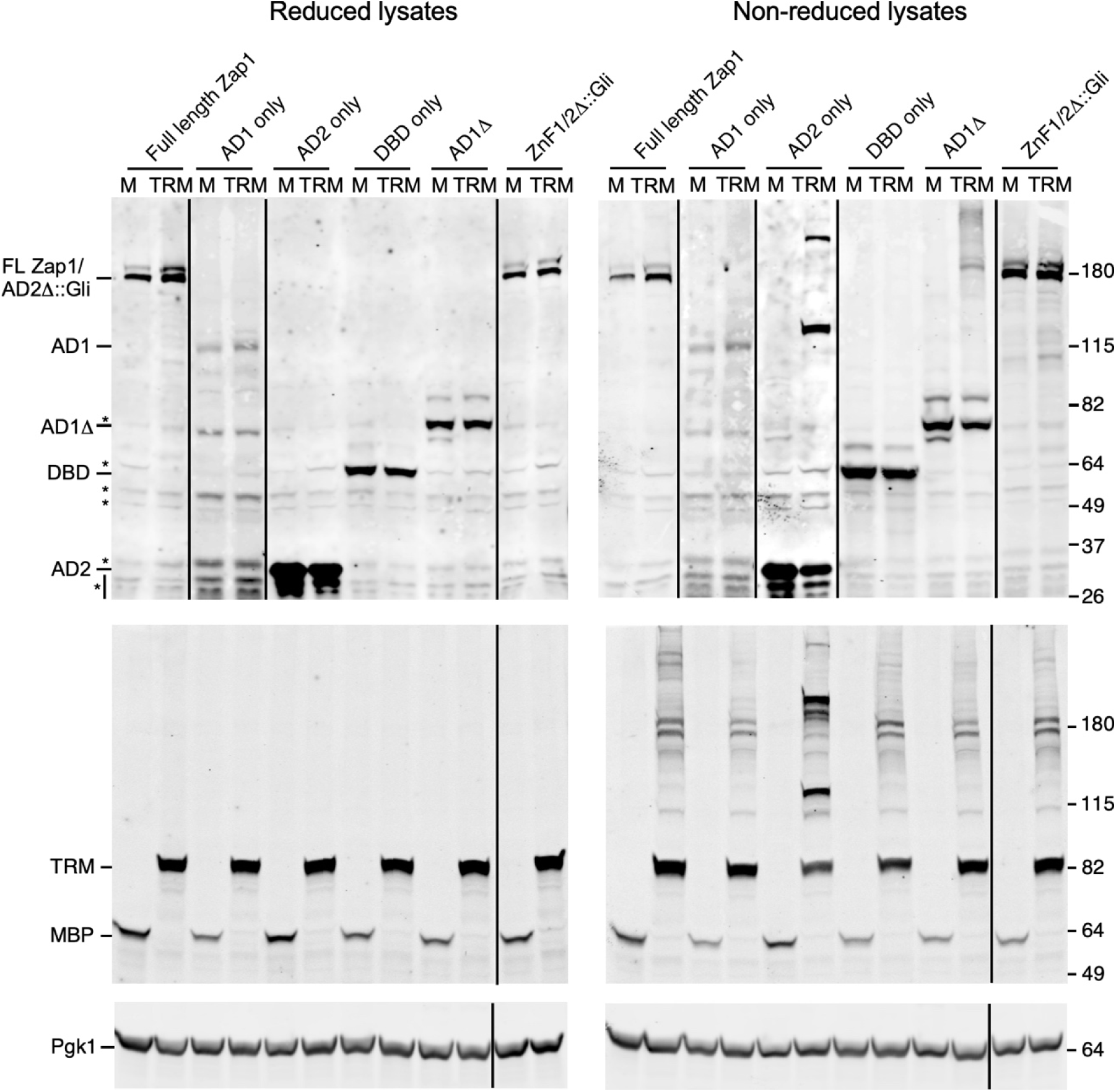
Immunoblot analysis of lysate fractions for Zap1 deletion analysis. Lysates of ZnR wild-type cells expressing MBP and the indicated Zap1 derivative, or *tsa1Δ* mutant cells expressing Tsa1^C171S^-MBP (TRM) and Zap1 derivatives were subject to SDS-PAGE and immunobotting to detect MBP (*middle* panels), myc-tagged Zap1 (*top* panels), or Pgk1 as loading and transfer control (*bottom* panels). Samples were reduced before loading (*left* panels) or untreated (*right* panels). Black lines indicate removal of superfluous lanes. For AD1 only lanes, image contrast was adjusted to more clearly show the low abundance AD1 band. Major non-specific bands generated by the anti-myc antibody are indicated with *asterisks* (*left top* panel).

**Supplemental Figure S11.**
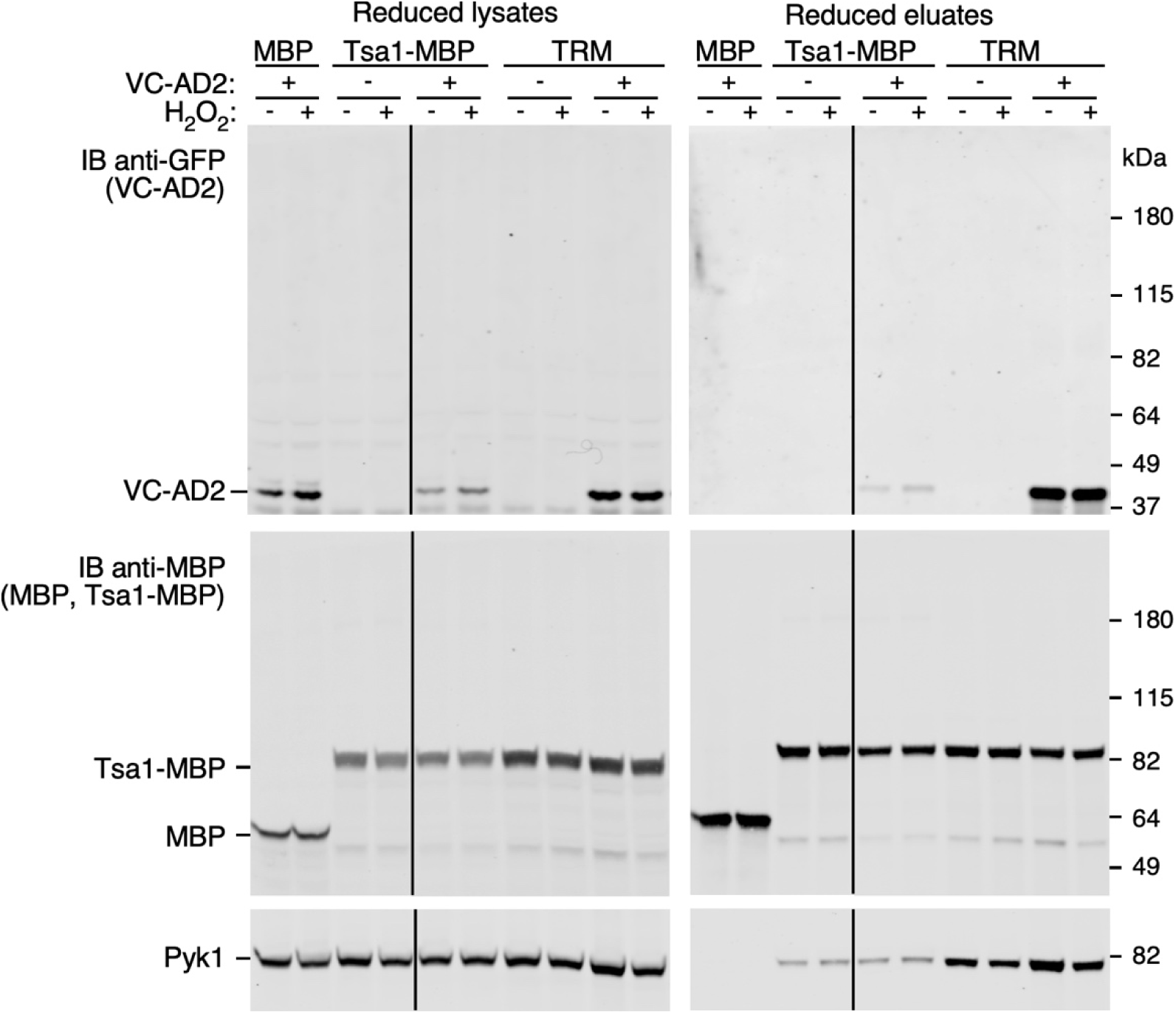
Wild-type Tsa1 oxidatively crosslinks to Zap1 AD2 *in vivo*. Samples were prepared as described in the legend to **Fig. 11**, and were reduced with DTT prior to SDS-PAGE and immunoblotting with antibodies to GFP (*top* panels), MBP (*middle* panels) and Pyk1 (*bottom* panels). Bands running below Tsa1-MBP (*middle* panels) are minor degradation products. Vertical lines indicate removal of superfluous lanes.

